# Cellular metabolic reprogramming controls sugar appetite

**DOI:** 10.1101/704783

**Authors:** Zita Carvalho-Santos, Rita Cardoso Figueiredo, Ana Paula Elias, Carlos Ribeiro

## Abstract

Cellular metabolic reprogramming is an important mechanism by which cells rewire their metabolism to promote proliferation and cell growth. This process has been mostly studied in the context of tumorigenesis and less is known about its relevance for non-pathological processes and how it affects whole animal physiology. Here, we show that *Drosophila* female germline cells reprogram their carbohydrate metabolism, upregulating the pentose phosphate pathway (PPP) to produce eggs. Strikingly, this cellular reprogramming strongly impacts nutrient preferences. PPP activity in the germline specifically increases the animal’s appetite for sugar, the key nutrient fueling this metabolic pathway. We furthermore provide functional evidence that the germline alters sugar appetite by regulating the expression of the fat body secreted satiety factor *fit*. The cellular metabolic program of a small set of cells is therefore able to increase the animal’s preference for specific nutrients through inter-organ communication to promote specific metabolic and cellular outcomes.

## Introduction

Within organisms, different cellular populations can have fundamentally different metabolic needs. Over the last decade, there has been an increased appreciation that cells can undergo an orchestrated reprogramming of their metabolic capacities, not to react to specific metabolic challenges, but to acquire new cellular and biological functions. The most prominent example of such reprogramming, originally described by Otto Warburg, is the rewiring of metabolism observed in tumor cells (Pavlova and Thompson, 2016; Vander Heiden et al., 2009; Warburg, 1956; Warburg et al., 1926). The “Warburg effect” is characterized by an increase in aerobic glycolysis and a concomitant reduced reliance of cells on oxidative phosphorylation (DeBerardinis and Chandel, 2016; Pavlova and Thompson, 2016; Vander Heiden et al., 2009). While mostly discussed in the context of pathological proliferative states, it is clear that such reprogramming also occurs in physiological settings. This is best appreciated in the context of development, where the metabolic program of cells is intimately linked to both their stemness as well as their differentiation potential (Shyh-Chang et al., 2013). In this context, metabolic reprogramming is thought not to be reactive but instead prewired to guide specific developmental outcomes. Animals are thus home to a multiplicity of cells with different metabolic identities. How organisms satisfy the differing and sometimes op-posing nutritional needs of such different cellular populations and, conversely, how cellular metabolic reprogramming affects whole-animal physiology and dietary choices has been little explored.

Originally, the “Warburg effect” was thought to be linked to the energy household of cells. However, more recently it has been proposed that the main advantage of this reprogramming is boosting the synthesis of essential macromolecular building blocks required during phases of high cellular demands such as proliferation (DeBerardinis et al., 2008b; Heiden and DeBerardinis, 2017; Lunt and Vander Heiden, 2011). Proliferating and growing cells as the ones found in tumors have a very high demand for metabolites such as nucleotides, amino acids, lipids, and redox potential, which they meet by channeling carbohydrates from glycolysis into the pentose phosphate pathway (PPP) (Lunt and Vander Heiden, 2011; Stincone et al., 2015). Accordingly, multiple studies in vertebrates and *Drosophila*, point to high dietary carbohydrate intake as promoting tumor growth (Goncalves et al., 2019a,b; Hirabayashi et al., 2013). The lack of *in vivo* whole animal models precludes a better understanding of the regulation and importance of PPP induction upon metabolic remodeling in physiological contexts.

The *Drosophila* female germline has served as a powerful, experimentally versatile model for discovering and dissecting many important cellular and developmental processes (Barton et al., 2016; Johnston and Ahringer, 2010; Lehmann, 2012). Oogenesis starts with the asymmetric division of a set of pluripotent germline stem cells (GSCs) followed by a set of tightly controlled, rapid cell divisions, and the maturation of the resulting egg chambers which contain the oocyte (Bastock and Johnston, 2008; de Cuevas et al., 1997; McLaughlin and Bratu, 2015; Slaidina and Lehmann, 2014). Given the importance of metabolism in determining both stemness and cell fate identity, recent work has started unraveling the importance of specific metabolic programs during *Drosophila* oogenesis (Sieber and Spradling, 2017). Most studies have focused on the remodeling of oxidative phosphorylation and its consequences on oogenesis. It has long been known that in late oocytes, mitochondria are remodeled and become quiescent (Cox and Spradling, 2003; Dumollard et al., 2007; Wallace and Selman, 1990). During the early stages of oogenesis, ATP synthase is thought to be required for the correct determination of the oocyte (Teixeira et al., 2015). Intriguingly this process seems to be independent of the function of the ATP synthase complex during oxidative phosphorylation. At later stages, remodeling of the mitochondria is thought to lead to a global shift in carbohydrate utilization and a resulting accumulation of glycogen (Sieber et al., 2016). While these studies have focused on the importance of mitochondrial remodeling, little is yet known regarding the other metabolic needs of the germline. Furthermore, the metabolic processes controlling the early stages of oogenesis and how they impact egg production remain poorly understood.

While the ability of proliferating cells to synthesize building blocks is now recognized as key to their function, nutrient uptake remains a key factor underlying all metabolic traits of cells. Accordingly, the *Drosophila* female germline is exquisitely nutrient sensitive. This has been best characterized for dietary proteins (mainly provided by dietary yeast) and amino acids (Carvalho-Santos and Ribeiro, 2018; Drummond-Barbosa and Spradling, 2001; Hsu and Drummond-Barbosa, 2009; Leitao-Goncalves et al., 2017; Piper et al., 2014; Søndergaard et al., 1995). Removal of these nutrients leads to a rapid and drastic reduction in egg production. While the requirement of dietary carbohydrates for oogenesis has been hardly explored, it is known that 65% of carbon in the germline is derived from dietary carbohydrates (Min et al., 2006). Furthermore, in *Drosophila*, sugars have been found to be mainly channeled into the PPP (Eisenreich et al., 2004) suggesting that this metabolic pathway could also play an important role in the germline. Dietary sugars therefore seem to be a critical source for metabolites during oogenesis, potentially through the PPP. Nevertheless, the importance of carbohydrate metabolism and the PPP for oogenesis remains to be assessed.

Animals are able to adapt their feeding behavior to meet their current nutritional needs. Many animals, including humans, do so by developing specific appetites, increasing their consumption from specific food sources in response to changes in various internal states including nutritional and mating states (Itskov and Ribeiro, 2013; Simpson et al., 2015; Simpson and Raubenheimer, 2012). Insects, for example, modulate food preferences to compensate for lack of dietary salts, amino acids or sucrose. They also increase the intake of certain nutrients in an anticipatory manner in response to mating (Corrales-Carvajal et al., 2016; Itskov et al., 2014; Leitao-Goncalves et al., 2017; Ribeiro and Dickson, 2010; Simpson et al., 2006; Trumper and Simpson, 1993; Walker et al., 2015). Changes in nutritional preferences are thought to be triggered by two mechanisms: the direct detection of changes in nutrient availability at the level of the central nervous system using neuronal nutrient sensing, and the reception of indirect nutritional information mediated by endocrine signals reporting the nutritional status of peripheral organs (Coll et al., 2007; Droujinine and Perrimon, 2016; Friedman and Halaas, 1998; Itskov and Ribeiro, 2013; Leopold and Perrimon, 2007; Pool and Scott, 2014; Williams and Elmquist, 2012). In *Drosophila* the female germline has been proposed to modulate food intake through ecdysone, controlling the increase in lipid accumulation in late-stage oocytes (Sieber and Spradling, 2015). It has however been shown that the germline and ecdysone do not play a role in increasing pro-tein appetite in response to amino acid deprivation or mating (Carvalho-Santos and Ribeiro, 2018; Ribeiro and Dickson, 2010; Walker et al., 2015). Whether the germline affects carbohydrate-specific appetites, and if so, how, is not known. In this study we describe two novel roles for sugar metabolism in the *Drosophila* female germline: promoting egg production and ensuring sugar intake. We demonstrate that both genetically ablating the ability of the germline to metabolize carbohydrates as well as dietary deprivation of this nutrient reduces egg production. The PPP plays a key role in mediating the impact of sugar metabolism on oogenesis. Genetically interfering with PPP activity in the ovaries severely reduces egg production. The ability of the germline to metabolize sugars through the PPP arises through the metabolic reprogramming of cells as they progress through oogenesis after differentiating from germline stem cells. By inducing the expression of the key carbohydrate metabolic enzyme *Hexokinase A* and enzymes of the PPP, these cells become competent to metabolize carbohydrates through this pathway, promoting egg production. We furthermore show that PPP activity in the germline induces a feed-forward increase in sugar appetite. Females without a germline or lacking PPP activity in this tissue show a drastic decrease in sugar appetite. This effect is specific for sugar feeding and relies on the increase in expression of the fat body secreted satiety factor Fit. *fit* mutants have high sugar appetite even when the germline is ablated. Our work highlights the importance of carbohydrate metabolic reprogramming for germline function, pinpoints the PPP as a key metabolic pathway required for egg production, identifies a novel feed-forward motif by which the metabolic identity of a small set of cells promotes the ingestion of the nutrient required for fueling this specific metabolic pathway, and provides functional evidence that this behavioral regulation relies on inter-organ communication between the germline and the fat body.

## Results

### Carbohydrate metabolism in the germline is required for egg production

Understanding the impact of nutrients and their metabolism on organismal function is a highly relevant but complex task. Carbohydrate metabolism has recently emerged as a key factor controlling cell proliferation and growth (Heiden and DeBerardinis, 2017; Lunt and Vander Heiden, 2011; Shyh-Chang et al., 2013). We therefore set to carefully dissect the effect of sugars on egg production and ovary physiology. In order to explore the specific impact of carbohydrates in an organ-specific manner, we decided to genetically interfere with the capacity of the germline to metabolize glucose. Hexokinases catalyze the initial step in the oxidative phosphorylation of hexoses (Fig. 1A). These enzymes are widely accepted to control glucose flux into different metabolic pathways (Stryer, 1995). In *Drosophila*, four different genes encode Hexokinases, giving rise to several isozymes. Of these, *Hexokinase A (HexA)* is thought to be the main isozyme expressed in the ovaries (Cavener, 1980; Chintapalli et al., 2007). We therefore specifically knocked down *HexA* in the female germline using the strong germline driver, *MTD-Gal4*, and assessed the resulting phenotypes in egg production and ovary morphology (Fig. 1B). Mated females with germline-specific *HexA* knockdown, displayed a dramatic decrease in the number of eggs laid in 24 hours when compared to control females (Fig. 1C). The inspection of egg chamber morphology within ovaries of these flies revealed readily observable abnormalities resulting from *HexA* knockdown (Fig. 1D and E). These included the presence of micronuclei which are suggestive of apoptosis.

**Figure 1.**
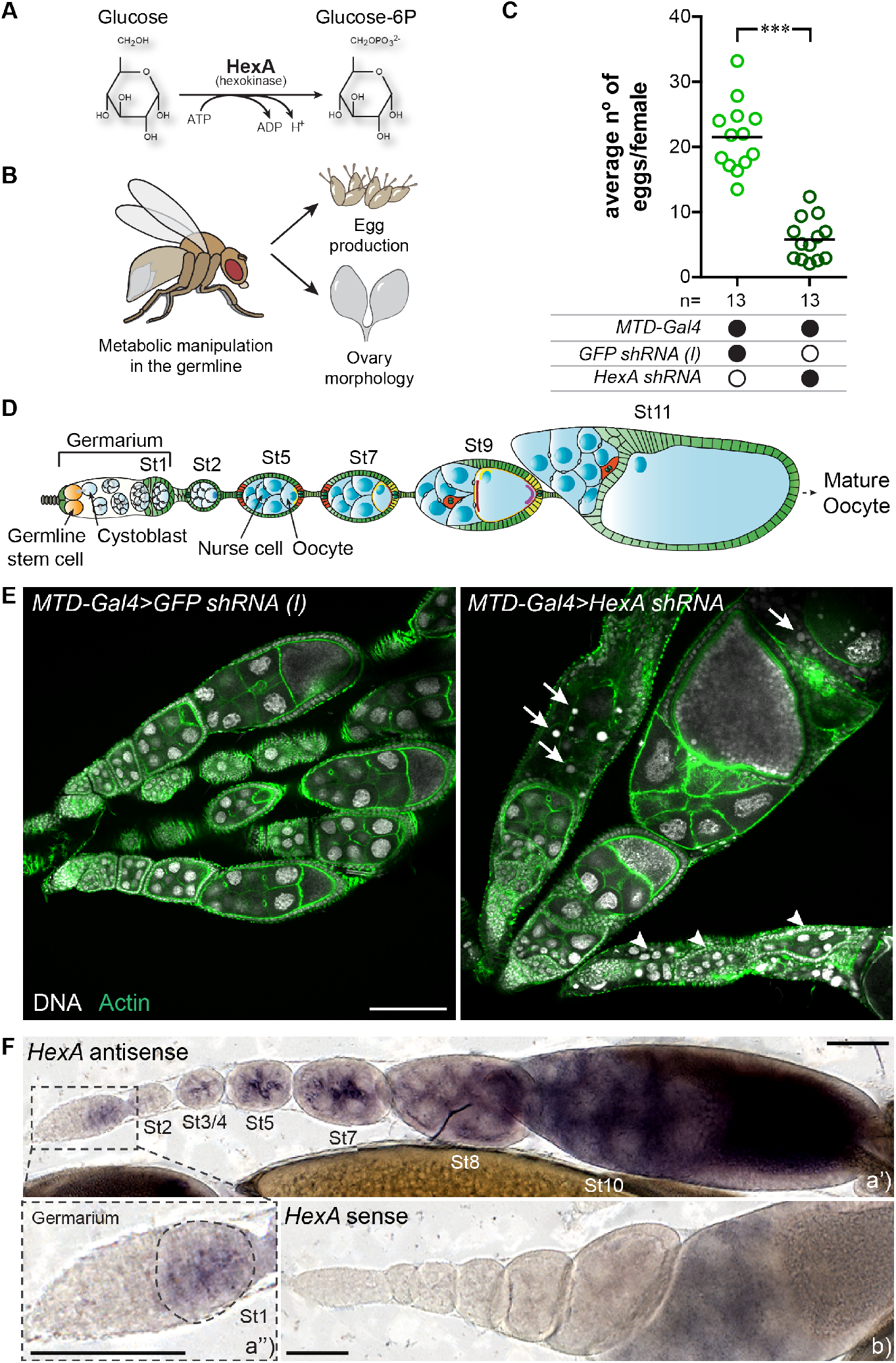
The germline undergoes a reprogramming of its carbohydrate metabolism which is required for egg production. (A) Schematic depicting the enzymatic reaction catalyzed by hexokinase, HexA. (B) The role of glucose uptake by the germline in egg laying and ovary morphology was assayed by knocking down the hexokinase *HexA* specifically in the germline. (C) Number of eggs laid per female in 24 h. *MTD-Gal4* driver was used to drive short hairpin RNAs specifically in the germline. A *GFP* knockdown line was used as a negative control. Black filled circles represent the presence and open black circles represent the absence of a particular transgene. Each colored circle in the plot represents eggs laid in single assays of 10-16 mated females (n = number of assays), with the line representing the mean. Statistical significance was tested using an unpaired t test. *** p < 0.001. (D) Schematic depicting a *Drosophila* ovariole (modified from Bastock and Johnston (2008)). Each ovary contains 22 ovarioles, composed of egg chambers in different developmental stages (St). The germarium, with the germline stem cells (GSCs) is localized at the most anterior tip. GSCs divide to produce cystoblasts which undergo four rounds of mitotic divisions. One of the 16 cystoblasts differentiates into the oocyte and the remaining develop into nurse cells. These cysts bud off of the germarium to form egg chambers which can be categorized into 14 different stages (St1- St11 depicted here) as they progress through oogenesis. (E) Representative ovariole morphology revealed by immunostaining of ovaries from females in which *HexA* was knocked down in the germline and the corresponding control. Arrows point to micronuclei and arrowheads to abnormal egg chambers. Green: actin, Gray: DNA. Scale = 100 μm. (F) Visualization of *HexA* mRNA expression in a representative ovariole using *in situ* hybridization. a’) *In situ* hybridization of ovaries using a *HexA* antisense probe. The dash-lined square represents a zoomed in view of the germarium (1.25x) (a”). b) *In situ* hybridization of ovaries using a *HexA* sense probe as a negative control. Scale = 100 μm. (C, E and F) Full genotypes of the flies used in these experiments can be found in Table S1.

### The female germline undergoes metabolic reprogramming

The metabolic program of different cellular populations is highly regulated, allowing them to adopt new and specific functions within the organism (Giese et al., 2019). When dysregulated, such metabolic changes can lead to highly aggressive pathologies such as cancer (Pavlova and Thompson, 2016; Sieber and Spradling, 2017). In order to analyze how carbohydrate metabolism is regulated in the germline, we visualized *HexA* expression using *in situ* hybridization and took advantage of the fact that the developmental progression during oogenesis is spatially organized in a linear fashion within the germline (Fig. 1D). In contrast to what would be expected for a housekeeping gene, the expression of *HexA* was not detected in all cellular populations of the germarium (Fig. 1F). We could not detect *HexA* mRNA in the most anterior part of the germarium, where the GSCs and cystoblasts are located (Fig. 1F a’ and a”). Expression of *HexA* becomes visible in the most posterior part of the germarium and as oogenesis progresses in both nurse cells and the oocyte (Fig. 1F a and a”). These data show that carbohydrate metabolism is not constitutively active throughout the germline but that the germline undergoes metabolic reprogramming when it transitions to more differentiated stages (Fig. 1F a’ and a”). Overall, our results indicate that the germline metabolizes dietary carbohydrates to generate eggs, a process that is critically mediated by *HexA*. These results explain the earlier reports showing that the female germline absorbs a high proportion of dietary carbohydrates, which in turn contribute to a large fraction of metabolites found in this organ (Min et al., 2006; O’Brien et al., 2008). Furthermore, carbohydrate metabolism is likely to be regulated at the transcriptional level, which could allow different cellular populations to use their new metabolic identity to support specific cellular and developmental functions.

### Dietary carbohydrates are required for egg production

If cellular carbohydrate metabolism is critical for oogenesis, dietary carbohydrate supply could also modulate egg production. We therefore decided to test if the sugar content of the diet impacts egg laying. A key challenge in nutritional research is the difficulty in manipulating specific nutrients when using natural foods. We therefore took advantage of a fully chemically defined fly diet (Piper et al., 2014, 2017), which allows us to precisely control the nutrient content of the diet. After 3 days on this diet, mated females were tested for egg laying and dissected for the analysis of their ovary morphology (Fig. 2A). We found that similarly to what had been described for amino acid deprivation, the acute deprivation of dietary carbohydrates resulted in a significant decrease in the number of eggs laid per female (Fig. 2B). Consistently, carbohydrate deprivation also led to a readily observable decrease in ovary size when compared to females kept on a full diet (Fig. 2C). These results show that in mated females, dietary carbohydrates are crucial for maintaining a high level of egg production. A reduction in ingested sugars is therefore likely to negatively impact the cellular carbohydrate flux in the germline. Maintaining an adequate supply of this important nutrient is therefore key for optimal reproductive output.

**Figure 2.**
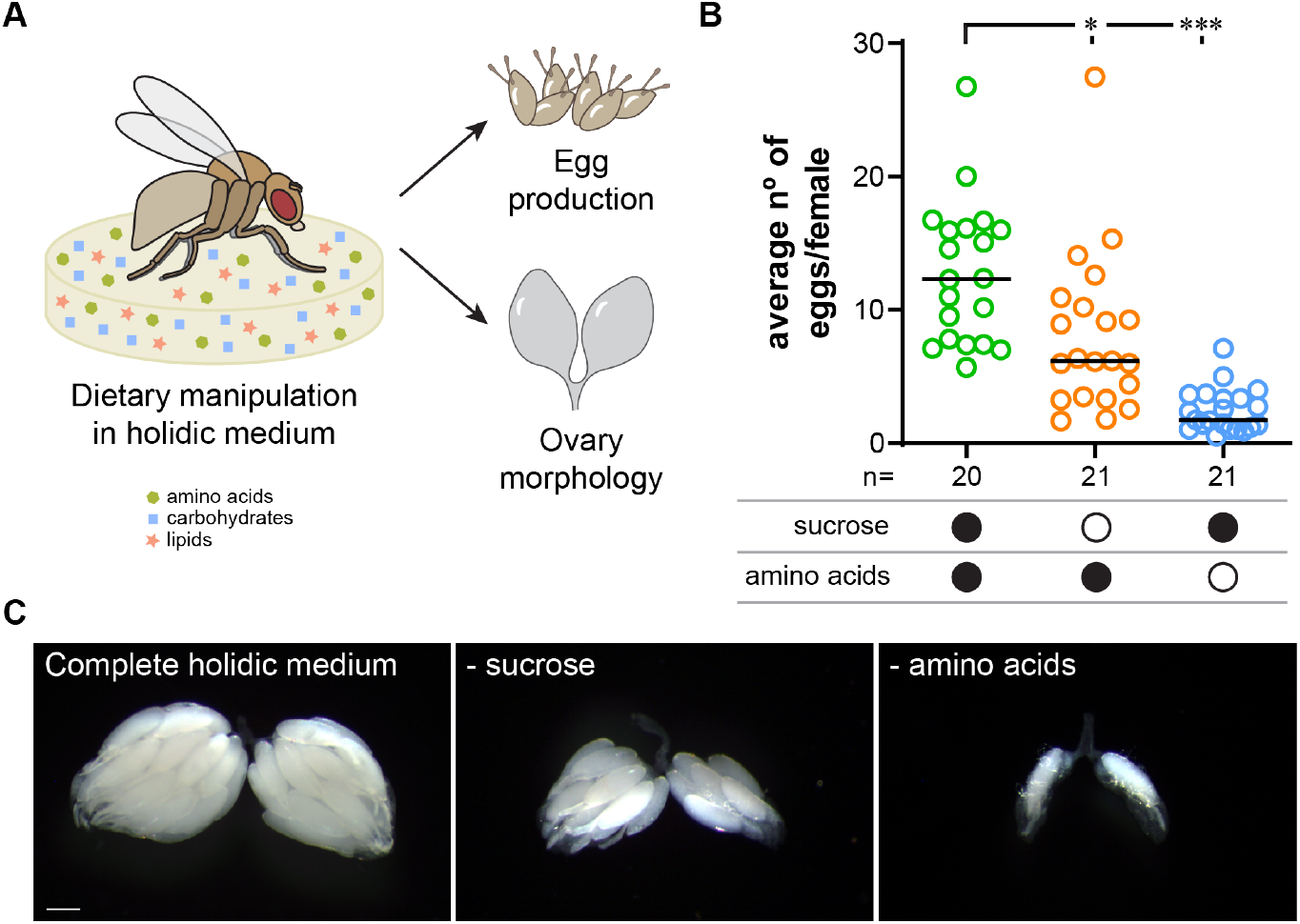
Dietary supply of sugars is required for egg production. (A) The holidic diet allows studying the impact of specific nutrients on fly physiology. Dietary manipulations using this medium were used to assess the role of sugars in egg laying and ovary morphology. (B) Average number of eggs laid per females in 24 h after feeding for 3 days on different holidic media. Black filled circles represent the presence and open black circles represent the absence of sucrose or amino acids in the holidic diet. Each colored circle represents the average number of eggs laid in single assays of 11-16 mated females (n = number of assays), with the line representing the median. Statistical significance was tested using the Kruskal-Wallis test followed by Dunn’s multiple comparison test. * p < 0.05, *** p < 0.001. (C) Ovary morphology of representative females fed for 3 days on full holidic diet and holidic diet lacking either sucrose or amino acids. Scale = 200 μm

### The germline modulates carbohydrate appetite

Food intake is the key process allowing animals to acquire all nutrients sustaining their metabolic requirements and supporting organ function. To maintain tissue nutritional homeostasis, animals adapt their foraging and feeding behaviors according to their current physiological needs (Corrales-Carvajal et al., 2016; Leitao-Goncalves et al., 2017; Ribeiro and Dickson, 2010; Simpson and Raubenheimer, 2012; Steck et al., 2018; Walker et al., 2015). Given that the germline requires a constant carbohydrate supply to sustain egg production, we decided to explore whether the germline could modulate sugar appetite. We tested this hypothesis by genetically ablating the germline and assaying the females for changes in feeding behavior (Fig. 3A). By overexpressing the transcription factor *bam*, which controls GSC differentiation in the germline, we induced a premature differentiation of the stem cells, resulting in females without a germline (Fig. 3B-E) (Ohlstein and McKearin, 1997). We tested these females for sugar appetite phenotypes using the flyPAD technology (Itskov et al., 2014). As expected females with a germline showed a clear increase in sugar appetite upon carbohydrate deprivation (Fig. S1A), which is best visualized by plotting the difference in sugar feeding between sugar deprived and fed flies (Fig. 3H). In contrast, this deprivation-induced increase in sugar feeding was abolished in females lacking a germline, which always showed a low level of sugar feeding (Fig. 3H and Fig. S1A). This phenotype is nutrient-specific as we never observed an alteration in amino acid deprivation-induced yeast appetite in these animals (Fig. 3I and Fig. S1B). Mating dramatically increases egg production and protein appetite via the Sex Peptide pathway (Ribeiro and Dickson, 2010). The modulation of sugar appetite by the germline is however not induced by mating, as virgin females lacking a germline also show a strong decrease in the appetite for carbohydrates (Fig. S1C). Finally, we also validated the reduction in sugar appetite induced by the ablation of the germline using a different behavioral assay (Ribeiro and Dickson, 2010) (Fig. S1D). Overall, our results show that, in contrast to protein appetite (Carvalho-Santos and Ribeiro, 2018; Ribeiro and Dickson, 2010), the germline strongly affects carbohydrate appetite. Stem cells are characterized by unique metabolic programs (Shyh-Chang et al., 2013). We therefore wondered if the observed sugar appetite phenotype could be specifically due to the ablation of the stem cells in the germline. To ablate the germline while keeping the GSCs, we knocked down *bam* using the same germline-specific driver. The resulting differentiation blockade produced ovaries composed solely ofa large number of GSCs (Fig. 3F-G). Instead of reacting to sugar deprivation by increasing sugar feeding, these females exhibited the same low sugar appetite as germline-ablated females, while showing an intact AA deprivation-induced yeast appetite (Fig. 3J-K and Fig. S1E-F). These results are consistent with our failure to detect *HexA* expression in GSCs (Fig. 1F) and demonstrate that while the germline is essential in driving sugar appetite, GSCs do not contribute to the modulation of sucrose appetite. The behavioral phenotype observed in germline-ablated females is consistent with the hypothesis that ovaries inform the central nervous system of their nutritional requirements to promote sugar appetite. We next explored whether glucose uptake by the germline also underlies the modulation of sucrose appetite. We knocked down *HexA* specifically in this tissue and tested the change in feeding behavior of the corresponding females upon carbohydrate deprivation. Consistent with all our previous results, these females show a strong and specific reduction in the drive to eat sucrose (Fig. 3L-M and Fig. S1G-H). Interestingly, while *HexA* knock-down germlines display morphological defects, these females still have ovaries (Fig. 1E). This suggests that it is the metabolic program of the germline rather than its presence that controls sugar appetite. Together, these results suggest that the metabolic program of a specific group of germline cells expressing *HexA* strongly impacts sugar feeding.

**Figure 3.**
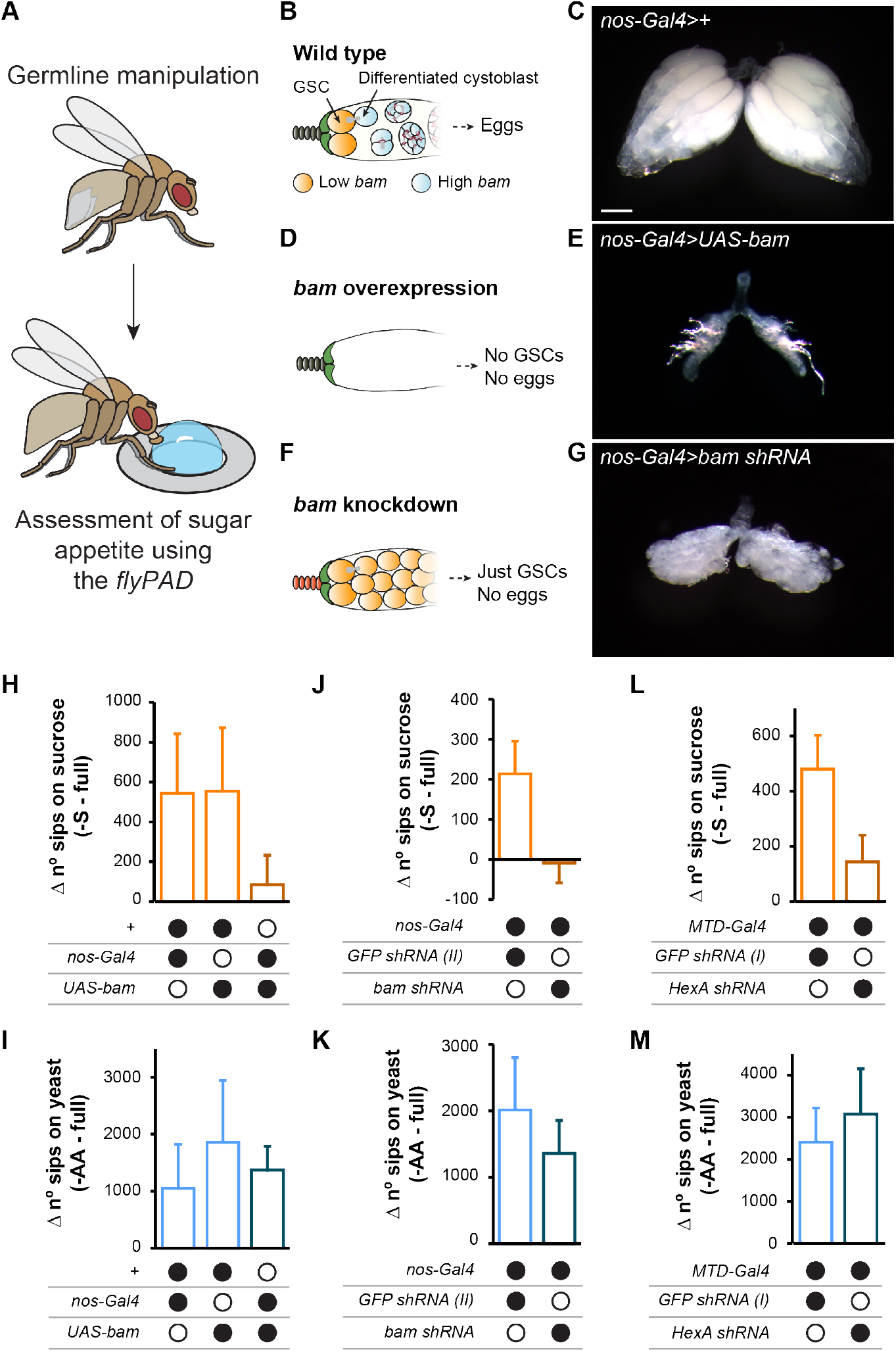
Carbohydrate metabolism in a subset of germline cells modulates sucrose appetite. (A) The role of the germline in nutrient appetite was assayed by full or partial ablation of the germline or knockdown of the hexokinase *HexA* in this tissue. (B, D and F) Schematic depicting the cellular effects of overexpressing or knocking down the transcription factor *bam* in the germline. (B) In wild type germaria, *bam* is transcriptionally repressed in the GSCs by Bone Morphogenetic Protein (BMP) signaling from the stem cell niche. Once GSCs divide, the daughter cell moves away from the niche leading to *bam* transcription and the differentiation into cystoblasts. (C) Ovary morphology from representative *wt* females. (D) *bam* overexpression in the germline leads to the premature differentiation of the GSCs and the rapid loss of germline cells and hence egg production. (E) Ovary morphology of representative females overexpressing *bam* specifically in the germline using *nos-Gal4* as a driver. (F) *bam* knockdown in the germline leads to a blockade in the differentiation into cystoblasts resulting in the accumulation of GSCs and the inability to produce eggs. (G) Ovary morphology of representative females with *bam* specifically knocked down in the germline using *nos-Gal4* as a driver. (H-M) Females in which the germline was fully (*nos-Gal4>UAS-bam*) or partially ablated (*nos-Gal4>bam shRNA*) or metabolically manipulated (M*TD-Gal4>HexA shRNA*) were assayed for an effect in nutrient choice using the flyPAD technology after 2 days on either a complete holidic medium or one lacking sucrose. Sucrose appetite is represented as the difference in sucrose feeding of flies maintained on holidic medium lacking sucrose (-S) vs full holidic medium (full) (H, J, and L). Yeast appetite is represented as the difference in feeding on yeast of flies maintained on holidic medium lacking amino acids (-AA) vs full holidic medium (full) (I, K, and M) (raw data in Fig. S1). Genotype matched *GFP* knockdown lines were used as negative controls in (J, K, L and M). (H-M) The columns represent the mean and the error bars show 95% confidence interval. Black filled circles represent the presence and open black circles represent the absence of a particular transgene. Full genotypes of the flies used in these experiments can be found in Table S1. Scale in C, E and G represents 200 μm.

### Despite their low drive to eat sugar, germline-ablated females are in a hungry state

The previous results can be easily explained if the ablation of the female germline leads to a drastic reduction in the utilization of existing energy reserves, and hence to an increase in resistance to starvation. To test if this could be the case, we measured glucose concentrations in the heads of germline-ablated females (Fig. 4A). As expected, sucrose deprivation led to a reduction in glucose in control animals. Germline ablation did not result in an increase in glucose in the head. If at all, these females showed a tendency towards a decrease in sugar concentration when compared to genetic background controls both in the fully fed and the carbohydrate-deprived situations (Fig. 4A). Furthermore, in contrast to their feeding behavior, germline-ablated females retain the ability to metabolically respond to dietary deprivation of sugar. They show a decrease in the concentration of glucose upon carbohydrate deprivation. Similarly, trehalose and fructose concentrations were also not increased in germline-ablated fly heads (Fig. S2). We further explored whether these females had increased available energy stores by carrying out a starvation resistance assay. In these experiments germline-ablated females did not show an increased resistance to starvation when compared to controls (Fig. 4B). This suggests that germline-ablated females do not have increased fat stores. Therefore the absence of sucrose appetite in females without a germline cannot be explained by the fact that these animals have higher energy reserves. Rather our results suggest that these females are in a metabolically “hyper-starved” state, which could result from their inability to increase sugar intake upon sugar deprivation.

**Figure 4.**
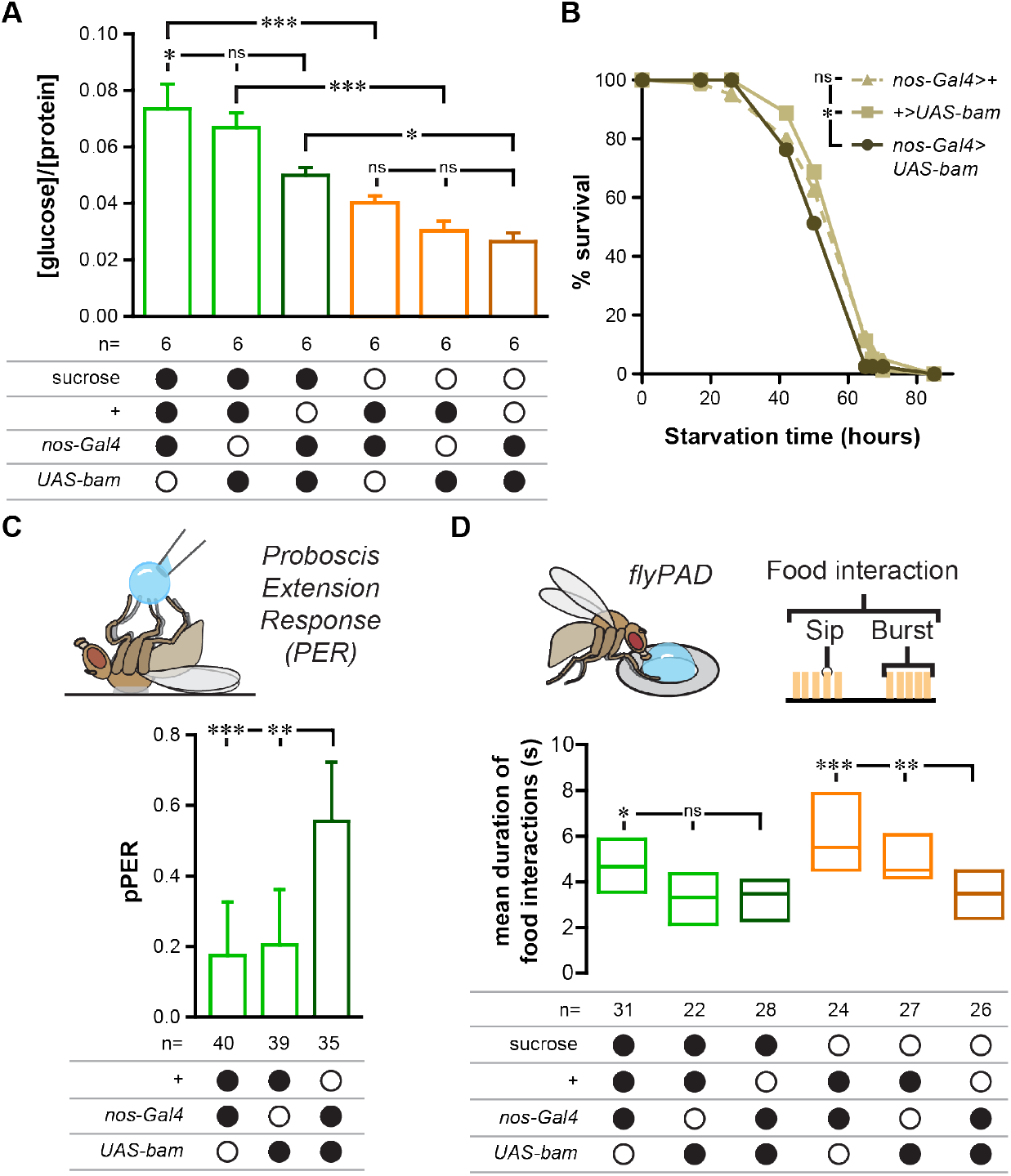
Germline-ablated females do not have increased available carbohydrates and are in a “hyper-starved” state. (A) Glucose measurements from heads of females reared for 2 days on full holidic medium (green) or holidic medium lacking sucrose (orange). Glucose concentrations were normalized to protein concentrations in the sample. The columns represent the mean and the error bars the standard error of the mean. n = total number of samples used per condition. Statistical significance was tested using an ordinary one-way ANOVA followed by Sidak’s multiple comparisons test. (B) Starvation curves of germline-ablated females and corresponding genetic background controls. Single flies were kept in tubes with water soaked paper and survival was scored every 12 h. n = 80. Statistical significance was tested using the Mantel-Cox test. (C) The probability of proboscis extension reflex (pPER) upon the presentation of a 25 mM sucrose solution to the tarsi was calculated for females maintained on a complete medium. The error bars show 95% confidence interval. Statistical significance was tested using Fisher’s exact test. (D) The mean duration of interactions with sucrose (s) was measured using the flyPAD setup using females maintained for 2 days on either a full holidic medium (green) or holidic medium lacking sucrose (orange). Boxes represent median with upper/lower quartiles. Statistical significance was tested using the Kruskal-Wallis test followed by Dunn’s multiple comparison test. (C and D) n = number of flies assayed per condition. (A-D) Black filled circles represent the presence and open black circles represent the absence of a particular nutrient in the holidic medium or of a given transgene. Full genotypes of the flies used in these experiments can be found in Table S1. ns p ≥ 0.05, * p < 0.05, ** p < 0,01, *** p < 0.001.μm.

### Females without a germline cannot sustain sugar feeding

The probability of initiating feeding is dependent on the detection of food by the gustatory system. Nutrient deprivation increases the sensitivity of gustatory neurons and hence the probability of proboscis extension (Inagaki et al., 2012; Steck et al., 2018). To test if the lack of the germline completely suppressed the ability of the female to react behaviorally to sugar deprivation, we specifically tested their drive to initiate feeding using the proboscis extension response (PER) assay. We calculated the PER of germline-ablated flies and corresponding genetic controls upon the presentation of a sucrose solution to the tarsal gustatory neurons (Fig. 4C). Despite their inability to increase sucrose feeding when carbohydrate deprived, fully-fed germline-ablated females displayed an increased PER to sugar when compared to controls (Fig. 4C). These data support our conclu-sion that these females are in a “hyper-starved” and hence carbohydrate-deficient state. But how can the overall lack of sugar feeding in these females be explained? Food intake is controlled by two opposite processes which either drive the animal to eat (hunger) or reduce its drive to forage and ingest food (satiation) (Dethier, 1976). The analysis of the different behavioral parameters generated by the flyPAD allows the behavioral separation of these two processes (Itskov et al., 2014). Indeed, a close analysis of the different feeding parameters generated by the flyPAD revealed that in control animals, upon sugar deprivation, females increased the time spent interacting with a sugar food spot (Fig. 4D). This behavioral phenotype is suggestive of a decrease in a satiation signal. Females without a germline however prematurely stop interacting with food. This suggests that while females lacking a germline have a strong drive to initiate feeding, sugar deprivation does not lead to a decrease in satiation. Collectively, our data suggest that a simple reallocation of resources from the germline to storage tissues does not explain the decrease in sucrose appetite observed in the germline-ablated flies. Instead, the hunger signal induced by dietary carbohydrate deprivation appears to be overruled by a dominant satiety signal that is active when the germline is absent.

### The PPP activity in the germline is essential for ovary function

Our data clearly show that dietary carbohydrates, and specifically sugar supply to the germline, is required for egg production (Fig. 1 and 2). This prompts the question as to which metabolic pathways in the germline utilize the ingested carbohydrates to support oogenesis. After phosphorylation by Hexokinase, glucose enters a variety of metabolic routes, the most prominent being glycolysis and the pentose phosphate pathway (Fig. 5A). The importance of these two pathways in driving cell proliferation in healthy and pathological states has nowadays been well documented in different organisms (Lunt and Vander Heiden, 2011; Sieber and Spradling, 2017; Stincone et al., 2015; Vander Heiden et al., 2009). Importantly, how metabolites flow through these two pathways has been associated with fundamentally different cellular proliferation and differentiation outcomes(Shyh-Chang et al., 2013). We therefore set out to characterize how metabolic pathways downstream of HexA in the female fly germline affect oogenesis. Classic work by Warburg has proposed that aerobic glycolysis is a key determinant of cell proliferation (Vander Heiden et al., 2009). To probe a possible involvement of glycolysis in oogenesis, we assessed egg laying in females in whose germline we knocked down three different enzymes of this pathway. In contrast to *HexA*, knockdown of *Pgi*, *Pfk* or *Pyk* in the germline did not lead to a decrease in egg production when compared to the corresponding genetic background controls (Fig. 5B). Given the high demand for building blocks in highly proliferating cells, the importance of glucose for these cells cannot be solely explained by their energy demands (DeBerardinis et al., 2008b; Lunt and Vander Heiden, 2011; Vander Heiden et al., 2009). This demand is thought to be met mainly by the synthesis of building blocks and generation of redox potential by the PPP (Fig. 5A). We therefore decided to knock down two enzymes of this pathway, *Zw* and *Pgd*, specifically in the germline to test the importance of this branch of carbohydrate metabolism in egg production. In contrast to the glycolysis genes, these manipulations led to a dramatic decrease in egg production when compared to the corresponding genetic controls (Fig. 5C). The effect was of a similar magnitude as the egg laying phenotype observed in *HexA* knockdown females (Fig. 1C). Given that knocking down different enzymes in the same pathway leads to the same phenotype, it is extremely unlikely that the observed phenotypes are due to off-target effects of the RNAi. Furthermore, our data is in line with the long known observation that mutants in purine biosynthesis, which is downstream of the PPP, show female sterility (Malmanche and Clark, 2004). The expression pattern of *HexA* in the germline suggests that this tissue undergoes a metabolic reprogramming to sustain high egg production. Indeed, *insitu* hybridization for *Pgd* mRNA localization in the germline revealed an identical expression pattern for this key PPP enzyme, strongly suggesting that the same reprogramming happens at the level of the PPP (Fig. 5D). We found that similarly to *HexA* (Fig. 1F), *Pgd* is only detectable in the most posterior part of the germarium and as oogenesis progresses in both nurse cells and oocyte (Fig. 5D a’ and a”). Overall, these results show that after GSC and cystoblast divisions, germline cells undergo a metabolic reprogramming, during which they transcriptionally induce the PPP, a key metabolic pathway important for egg production.

**Figure 5.**
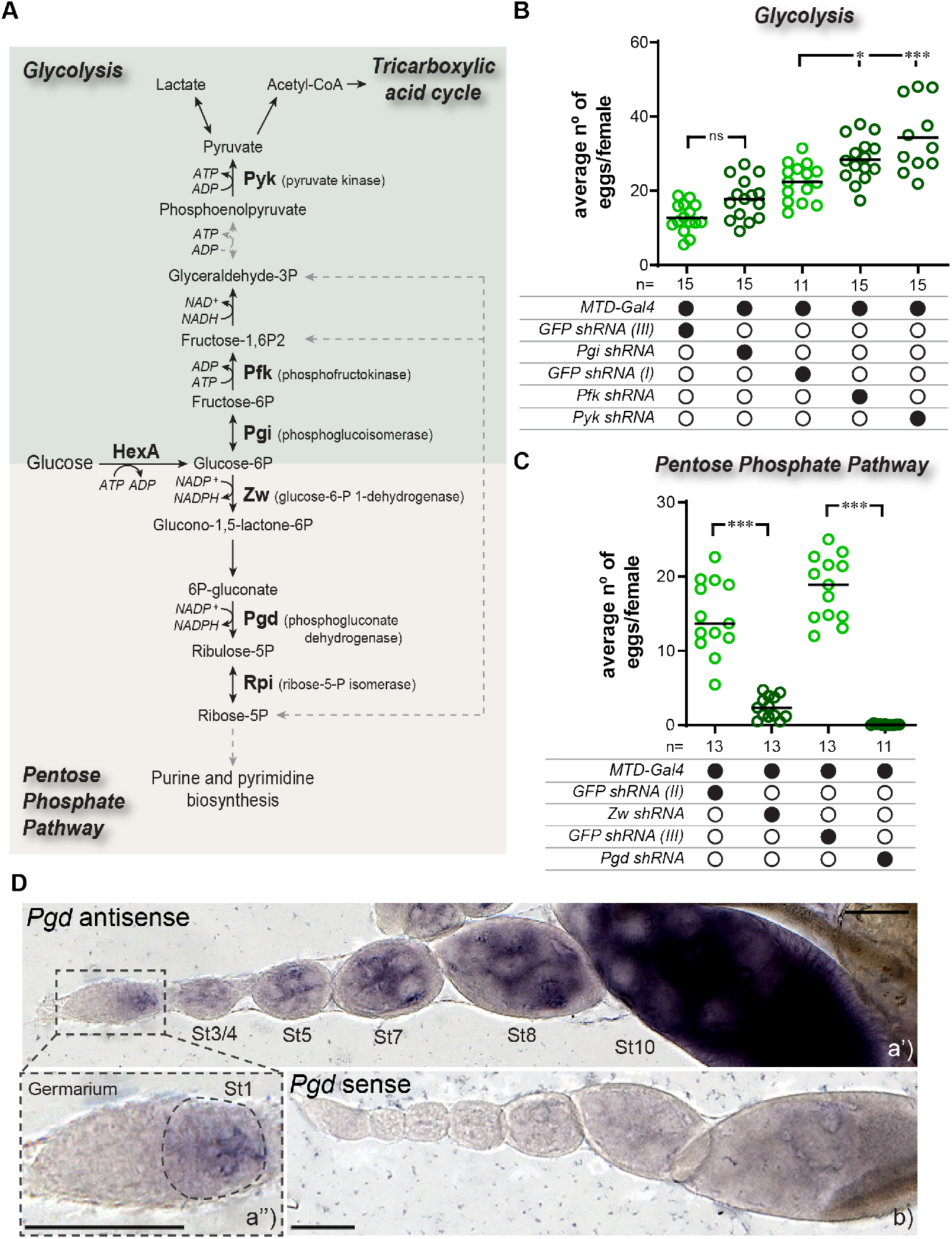
The activity of the pentose phosphate pathway in the germline is required for egg production. (A) Simplified schematic of the enzymatic reactions encompassing the two core metabolic pathways downstream of glucose phosphorylation by hexokinase, Glycolysis (light green) and the Pentose Phosphate Pathway (PPP) (light brown). Metabolic steps represented in dashed grey lines include more than one enzymatic reaction. (B and C) Average number of eggs laid per female in 24h. *MTD-Gal4* driver was used to drive short hairpin RNAs specifically in the germline. *GFP* knockdown lines were used as negative controls. Black filled circles represent the presence and open black circles represent the absence of a particular transgene. Each colored circle in the plots represents the average number of eggs laid in single assays of 13-17 mated females (n = number of assays), with the line representing the median. Statistical significance was tested using an ordinary one-way ANOVA followed by Sidak’s multiple comparisons test (B) and Mann-Whitney test (C). ns p ≥ 0.05, * p < 0.05, *** p < 0.001. (D) Visualization of *Pgd* mRNA expression in a representative ovariole using *in situ* hybridization a’) *In situ* hybridization of ovaries of control flies using a *Pgd* antisense probe. The dash-lined square represents a zoomed in view of the germarium (1,25x) (a”). b) *In situ* hybridization of ovaries of control flies using a *Pgd* sense probe as a negative control. St – Stage. Scale = 100 μm. Full genotypes of the flies used in these experiments can be found in Table S1.

### The PPP activity in the germline modulates sugar appetite

Our data suggest that the germline, and more specifically *HexA* activity in this tissue, promotes sugar appetite (Fig. 3). Moreover, the induction of the PPP during oogenesis is required for egg production (Fig. 5). We next investigated whether this pathway also modulates sugar appetite. We knocked down *Pgd* and *Zw* specifically in the germline and tested these females for changes in sugar appetite. Impairing the activity of the PPP by knocking down *Zw* or *Pgd* in the germline led to a dramatic decrease in the appetite for sucrose when compared to the corresponding genetic controls (Fig. 6A and Fig. S3A). We confirmed these results using a *Pgd*, *Zw* double mutant which has been previously characterized to almost completely abolish the metabolic flux through the PPP (Gvozdev et al., 1976; Hughes and Lucchesi, 1977) (Fig. 6B and Fig. S3B). Similarly to what we found for other germline manipulations, neither females with a germline knock-down of *Pgd*, *Zw* nor the double mutants showed defects in amino acid deprivation-induced yeast appetite (Fig. S3C-D). Collectively, these results show that the PPP metabolic program in the germline is not only crucial for egg production but also to promote sugar appetite. This links the metabolic activity of this pathway in the ovaries to the regulation of the intake of carbohydrates fueling it, resulting in the maintenance of a high reproductive output.

**Figure 6.**
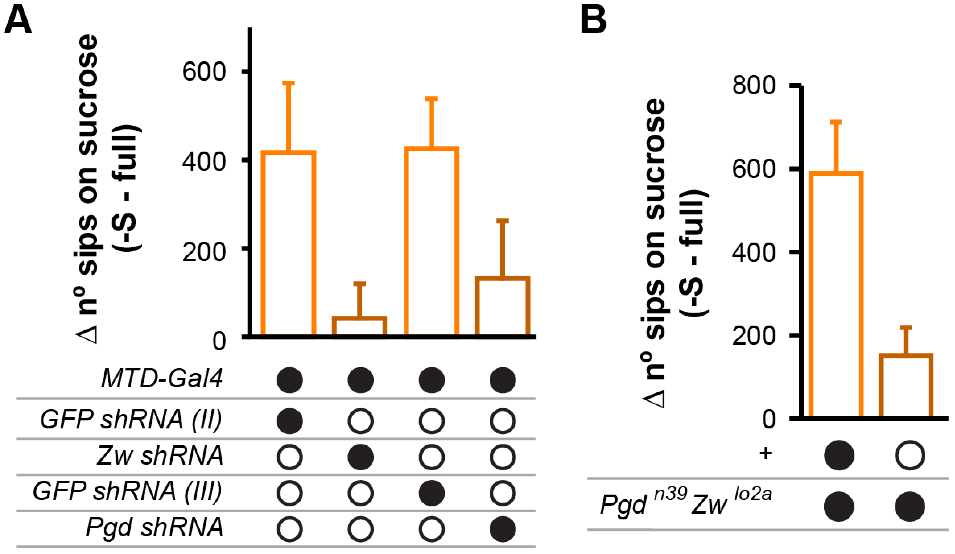
The pentose phosphate pathway activity in the germline modulates sucrose appetite. (A-B) Flies were assayed for feeding behavior after 2 days on either a complete holidic medium (full) or one lacking sucrose (-S). Sucrose appetite is represented as the difference in sucrose feeding of flies maintained on holidic medium lacking sucrose *vs* full holidic medium (raw data in Fig. S3). (A) The *MTD-Gal4* driver was used to drive short hairpin RNAs specifically in the germline and matching *GFP* knockdown lines were used as a negative control. (B) A *Pgd*, *Zw* double-mutant was used to interfere with the PPP pathway. The columns represent the mean and the error bars show 95% confidence interval. Black filled circles represent the presence and open black circles represent the absence of a particular transgene or a mutation in homozygozity. Full genotypes of the flies used in these experiments can be found in Table S1.

### The fat body secreted satiety factor Fit mediates the germline regulation of sugar appetite

In both vertebrates and invertebrates, the systemic adaptation of physiology and behavior to the availability of nutrients relies on inter-organ communication (Droujinine and Perrimon, 2016; Williams and Elmquist, 2012). In vertebrates the adipose tissue and the liver play pivotal roles in this crosstalk (Williams and Elmquist, 2012). In invertebrates the fat body fulfills a similar role as a key coordinator of nutritional homeostatic responses (Leopold and Perrimon, 2007). We therefore reasoned that secreted factors expressed in the fat body of females could be valid candidates for mediating the germline effect on sugar cravings. female-specific independent of transformer (*fit*) encodes a secreted peptide which is expressed in the fat body tissue surrounding the brain of females (Fujii and Amrein, 2002). Moreover, the nutritional state of the animal regulates *fit* expression (Fujikawa et al., 2009). We therefore hypothesized that Fit is involved in communicating the metabolic state of the female germline to the brain. We started by testing this hypothesis by assessing if *fit* expression is regulated by the germline and its metabolic state. Indeed, we found that while in control females which have been sugar deprived, *fit* is expressed at very low levels, both germline ablation, as well as germline knockdown of HexA, led to a very clear increase in *fit* expression (Fig. 7A and S4A). This effect is in agreement with earlier observations showing that progeny of *Tudor* mutant females, lacking a germline, also show a drastic increase in *fit* expression (Parisi et al., 2010). The regulation of *fit* therefore supports the hypothesis that the fat body senses the metabolic state of the germline and secretes a satiety factor that modulates sugar intake. To functionally test whether the increased expression of *fit* in germline-ablated flies underlies their decreased sugar appetite, we assessed if removal of *fit* from these flies would restore increased sugar intake. We combined a *fit* null mutation (Sun et al., 2017) with a transgene that allows the expression of *bam* under the control of a heat shock promoter (Fig. 7B). As observed in control animals, female *fit* mutants with an intact germline responded to sugar deprivation by increasing their carbohydrate consumption (Fig. 7C and S4B). This result is consistent with the observation that in control sugar-deprived females, *fit* is already hardly expressed (Fig. 7A and S4A). Germline ablation in a heterozygous *fit* mutant background using a heat shock treatment clearly reduced the feeding on sugar as observed in other germline-ablated females (Fig. 7C and S4B, Fig. 3H and S1A). Strikingly however, females with an ablated germline and homozygous *fit* mutant background, showed a rescue of the appetite for sucrose to a level comparable to females with a germline (Fig. 7C). These results suggest that similarly to what has been shown for protein intake, Fit acts as a satiety factor to suppress sucrose appetite (Sun et al., 2017). They also nicely explain our earlier observations that the germline does not control sugar feeding initiation but feeding maintenance, a behavioral pattern suggesting the involvement of a satiation factor (Fig. 4D). Our data suggest that Fit controls satiety by integrating the metabolic activity of the female germline to fine tune the intake of carbohydrates. By acting as a multiorgan relay, the fat body participates in matching the intake of carbohydrates to the metabolic needs of the PPP in the germline, promoting the continuous availability of sugars required for egg production (Fig. 7D). This anticipatory feed-forward mechanism could be just one example of a more general strategy by which metabolically distinct cell populations communicate their specific needs to the brain to ensure the intake of nutrients vital to their metabolic needs.

**Figure 7.**
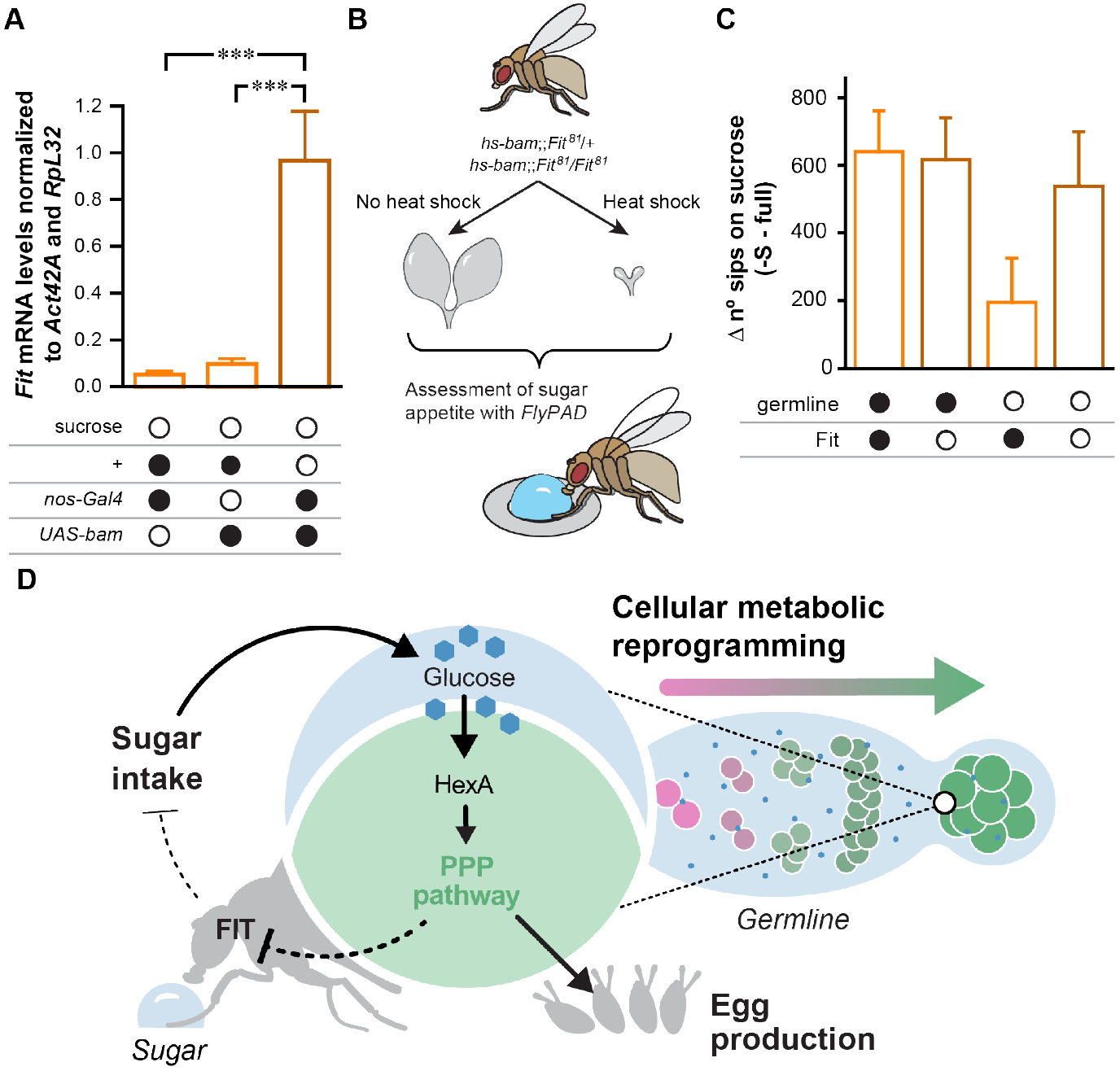
The germline modulates sugar appetite by regulating the expression levels of the fat body secreted satiety peptide *fit*. (A) *fit* mRNA levels were measured in whole, mated female flies fed on holidic medium lacking sucrose and normalized to two internal controls (*Actin 42A* and *RpL32*). Black filled circles represent the presence and open black circles represent the absence of a particular nutrient in the holidic medium or of a transgene. The columns represent the mean and the error bars the standard error of the mean. Statistical significance was tested using an ordinary one-way ANOVA followed by Sidak’s multiple comparisons. (B) Females heterozygous or homozygous for a *fit* null mutation and carrying a *hs-bam* transgene were heat shocked during development to generate germline-ablated animals. Control animals with a germline were generated by skipping the heat treatment. These flies were then assayed for their sucrose appetite. (C) Flies were assayed for an effect in nutrient feeding using the flyPAD technology after 2 days on either a complete holidic medium or one lacking sucrose. Sucrose appetite is represented as the difference in sucrose feeding of flies maintained on holidic medium lacking sucrose (-S) vs full holidic medium (full) (raw data in Fig. S4). The columns represent the mean and the error bars show 95% confidence interval. Black filled circles represent the presence and open black circles represent the absence of the germline or the *fit* gene. Full genotypes of the flies used in these experiments can be found in Table S1. ns p ≥ 0.05, * p < 0.05, *** p < 0.001. (D) Model depicting how cellular metabolic reprogramming impacts organ function and carbohydrate appetite. As oogenesis progresses, germline cells undergo metabolic reprograming (right) and dramatically increase the expression of carbohydrate metabolism and PPP genes (center). This directs dietary carbohydrates into the PPP pathway, which is essential for egg production (center). Carbohydrate flux through the PPP pathway increases sugar appetite by suppressing the expression of the satiety peptide Fit in the head fat body (left). The resulting increase in sugar appetite sustains the glucose flux through the PPP in the germline and hence egg production in a feed-forward manner.

## Discussion

Cellular metabolic reprogramming is an important biological process by which cells rewire their metabolism to promote cell proliferation, cell growth, and specific developmental outcomes (DeBerardinis et al., 2008a; Lunt and Vander Heiden, 2011; Shyh-Chang et al., 2013; Sieber and Spradling, 2017; Vander Heiden et al., 2009). This process has been mostly studied in pathological or *ex vivo* conditions. We identified a new case of cellular metabolic reprogramming in which cells in the female reproductive organ of *Drosophila* rewire their metabolism allowing them to utilize the pentose phosphate pathway to generate eggs. We also show that this new metabolic program profoundly impacts whole organism physiology. Gonadal carbohydrate metabolism alters the expression of the satiety factor Fit. Fit is known to be secreted by the head fat body surrounding the brain of the adult female fly, on which this peptide acts to influence feeding. We find that by dramatically reducing the expression of the satiety factor *fit*, carbohydrate flux in the female gonads specifically increases sugar intake. As dietary sugars are key for maintaining a high reproductive rate, this feed-forward regulatory loop ensures the adequate provisioning of carbohydrates to fuel the PPP and hence reproduction.

The Warburg effect is widely regarded as the canonical example of metabolic reprogramming (Vander Heiden et al., 2009; Warburg et al., 1926). This alteration of cellular metabolism in tumors is characterized by an increase in aerobic glycolysis and a concomitant production of lactate. Metabolically reprogrammed cells also display a dramatically increased consumption of carbohydrates (Gambhir, 2002). The Warburg effect was long thought to be intimately linked to the energetic demands of cells. But over the last year, the importance of the Warburg effect for the generation of building blocks has emerged as a crucial benefit of cellular metabolic reprogramming (DeBerardinis et al., 2008b; Heiden and DeBerardinis, 2017; Lunt and Vander Heiden, 2011). Proliferating cells have a high demand for nucleotides, fatty acids, amino acids, and redox potential. These are all metabolic products generated by the PPP from carbohydrates (Stincone et al., 2015). Indeed, aggressive tumors are known to increase the flux of carbohydrates through the PPP by downregulating their flux through glycolysis (DeBerardinis et al., 2008b). In this context it is interesting to note that in flies in which we interfered with the expression of glycolytic enzymes, egg laying increased (Fig. 5B). These data suggest that similarly to what happens in some tumors, reducing the flux through glycolysis leads to an increase in PPP flux and hence the production of eggs. Overall our data strongly suggest that the flux through the PPP is rate-limiting for egg production. Our work therefore adds to the body of work linking the Warburg effect to the production of building blocks and redox potential through the PPP. Appropriately, Otto Warburg not only discovered the metabolic reprogramming phenomenon named after him but also led the efforts culminating in the identification of a key enzyme in the PPP (Warburg and Christian, 1936; Warburg et al., 1935). Almost 100 years after his seminal work, biologists are now stitching together his findings into a coherent picture, linking cellular metabolic reprogramming to the biosynthetic capacities of the PPP.

Our findings prompt the question of the extent to which cellular metabolic reprogramming could be relevant for reproduction in other organisms. While detailed molecular studies in vertebrates have not fully addressed this question in the context of the whole animal, experiments performed with ES cells as well as knowledge from in vitro fertilization clinical practice suggest that metabolic reprogramming also plays an important role in reproduction across phyla. It is nowadays widely appreciated that changes in metabolism play an important role in instructing specific developmental fates early in vertebrate development (Shyh-Chang et al., 2013). Most of these changes are linked to alterations in carbohydrate metabolism and have partially been linked to the utilization of the PPP. It is therefore very likely that during human reproduction, metabolic rewiring also plays an important role. If this is linked to changes in appetite and how potential cellular metabolic alterations are linked to whole animal physiology is unknown. What is clear is that in animals including humans, reproduction is intimately linked to changes in appetite and food preferences (Walker et al., 2017). Intriguingly, in women, energy expenditure increases by 10% during the luteal phase of the menstruation cycle (Webb, 1986) and multiple studies have reported increased consumption of carbohydrates during the premenstrual period (Bryant et al., 2006; Dye and Blundell, 1997) which has been linked to changes in sucrose thresholds (Than et al., 1994). The reported effects are however modest and are partially contested. Mechanistic studies addressing the importance of cellular metabolic reprogramming in the context of physiological processes such as reproduction and nutritional behavior are likely to bring more clarity and be a fertile area for future research.

Why is the PPP so important for oogenesis? Most likely it is not one specific metabolic product of this pathway which justifies the profound rewiring we observe in our study, but the full set of the produced building blocks and the redox potential essential for cell proliferation and cell growth. But nucleotides might be key to at least partially understand the importance of the cellular biosynthesis of building blocks for reproduction and development. It has long been known that in *Drosophila* mutants affecting purine synthesis downstream of the PPP result in female sterility (Malmanche and Clark, 2004). Intuitively, one could argue that given the presence of nucleotides in the diet, the necessity for nucleotide biosynthesis in the germline should be minimal. But recent work has highlighted two important points which could explain this apparent contradiction. First, the pace of cell proliferation and cell growth often outpaces the capacity of cells to absorb building blocks. This makes proliferating and growing cells dependent on their biosynthetic capacity for nucleotides (Song et al., 2017) and might be especially important in endoreplicating cells such as the nurse cells. But more intriguingly, the exact levels of nucleotides during early embryogenesis is emerging as an important factor controlling early steps of embryonic development. Both low and high levels of nucleotides negatively affect morphogenetic processes in the early embryo (Djabrayan et al., 2019; Liu et al., 2019). Cells rely on the exquisitely precise regulatory control of ribonucleotide reductase enzymatic activity to ensure a precise regulation of nucleotide levels in the egg (Djabrayan et al., 2019; Song et al., 2017). It is therefore tempting to speculate that the germline induces PPP activity not only to ensure the availability of this critical nutrient but also to be able to regulate the levels of this metabolite in a diet-independent manner.

A key discovery of our study is the ability of metabolically reprogrammed cells to alter sugar appetite. This ensures the provisioning of these cells with an adequate supply of carbohydrates which then further fuels sugar appetite. We show that this change in appetite is mediated by inter-organ communication. Inter-organ communication plays a central role in relaying and coordinating the metabolic needs of organs and specific cellular populations to ensure homeostasis (Droujinine and Perrimon, 2016). These interactions can be mediated by secreted, dedicated signaling molecules (e.g. hormones), metabolites, or neuronal routes. The brain plays an important role in these interactions as decisions related to food intake are a primary means of regulating whole animal physiology. By directly influencing brain function, specific cellular populations can ensure that the feeding behavior of the whole organism meets the specialized metabolic need of small groups of cells. This is especially important when their needs deviate from the needs of the majority of cells in the animal. Reproduction is a state in which this situation is especially relevant. The generation of offspring imposes a high metabolic burden on the mother, both in terms of quantity and quality of nutrients. The increased needs for salt and proteins in reproducing females are met by anticipatory changes in salt- and protein-specific appetites (Walker et al., 2017). In vertebrates, these are induced by specific hormones. In the case of *Drosophila*, the mating state of the female is conveyed by specific neuronal pathways to the brain which control taste processing to generate nutrient specific appetites (Walker et al., 2015). Here we describe a novel strategy by which reproductive cells can alter feeding behavior to ensure that the animal ingests a diet rich in carbohydrates. Our data suggest that the carbohydrate flux through the PPP in the metabolically remodeled germline is sensed by the fat body leading to the transcriptional inhibition of the gene encoding the secreted peptide Fit. Indeed *fit* expression is upregulated in both females without a germline and females with germline-specific *HexA* knockdown. Importantly, *HexA* knockdown does not lead to germline ablation, indicating that it is not the absence of a germline per se which leads to changes in *fit* expression and sugar appetite. Fit is likely to act as a sugar satiety signal as flies without a germline and mutant for *fit* do not show a loss of sugar appetite. It is intriguing that we identify Fit as an important signal regulating sugar appetite, as a previous study has identified Fit as being regulated by the protein content of the diet in females and mediating the satiety effect of this nutrient (Sun et al., 2017). If one takes into account that dietary amino acids (AAs) have a profound impact on the female germline (Fig. 2), the regulation of *fit* by dietary proteins observed by Sun and colleagues could be explained by the indirect impact of AAs on the germline. In our model *fit* does not detect the presence of specific nutrients in a sexually dimorphic way as originally proposed, but would react to the activity of the germline and convey its metabolic state to regulate nutrient selection.

Many questions still remain to be addressed. Key will be the identification of the signal from the germline which is controlled by the carbohydrate flux through the PPP in this organ. The signal could be a hormone or a dedicated signaling protein. We have tested multiple likely candidates such as ecdysone (Carvalho-Santos and Ribeiro, 2018; Sieber and Spradling, 2015) and Dilp8 (Colombani et al., 2012; Garelli et al., 2012) and found no evidence for their involvement in sugar appetite (data not shown). An alternative possibility is that the signal is a metabolite produced by the flux of carbohydrates through the PPP which then directly acts on the fat body to regulate *fit* expression. Although identifying such factors has notoriously been difficult, a combination of genetic, transcriptomic, and metabolomic approaches should yield the identity of the mechanisms by which the germline acts on the head fat body to control *fit* expression. Another key question is how Fit acts to alter sugar appetite. Sun and colleagues proposed that Fit acts on insulin-secreting neurons to exert its satiety effects. While we have observed that germline ablation can lead to an alteration in Dilp2 and Dilp3 levels in female brains, these changes do not correlate with the observed alterations in feeding behavior (data not shown). The effects in Dilp levels are more readily explained by the observed changes in circulating sugars in these flies (Fig. 4). Identifying the molecular and circuit mechanisms by which Fit alters sugar appetite will be a key future avenue to understand how the metabolic state of the germline alters food preferences.

Anticipatory, feed-forward regulatory strategies are emerging as an indispensable principle ensuring nutritional and physiological homeostasis (Andermann and Lowell, 2017; Walker et al., 2017). Such anticipatory strategies guarantee a continuous supply of resources. Using this strategy, the animal circumvents the need for an error signal (induced by the lack of the nutrient) which triggers homeostatic compensation in pure feedback regulatory systems. We propose that we have identified a novel example of such a feed-forward regulatory strategy, which is especially relevant for understanding the impact of metabolic reprogramming. By coupling the flux of carbohydrates through the PPP to an increase in sugar appetite, metabolically reprogrammed cells ensure a continuous, uninterrupted availability of the key metabolic precursor fueling this metabolic pathway. By doing so, cellular metabolic reprogramming extends beyond the cellular level, leading to a whole-organism behavioral metabolic reprogramming. We propose that this strategy is likely not to be confined to the *Drosophila* female germline but could represent a generalizable regulatory strategy by which metabolically reprogrammed cells could alter physiology and feeding behavior across phyla. If this is the case, one of the most provocative predictions would be that metabolically reprogrammed tumorigenic cells could tap into such a feedforward regulatory loop to increase the appetite of the host for specific nutrients. Given that a high carbohydrate intake has been linked to an increase in tumor growth (Goncalves et al., 2019a,b; Hirabayashi et al., 2013) this would likely lead to a boost of the metabolic capacities of reprogrammed cells and hence proliferation and disease progression. Exploring to what extent metabolic programming rewires whole organism physiology and behavior, identifying the mechanisms underlying such systemic effects, and if this systemic rewiring can explain the progression of diseases relying on cellular metabolic reprogramming, promises to be a fruitful avenue for future research yielding novel insights into how animals maintain homeostasis.

## Supporting information

Supplemental data Santos et a

## ACKNOWLEDGEMENTS

We thank Ralph Neumüller, Dennis McKearin, Michael Buszczak, Yan Li, Rui Martinho, and Pedro Prudencio for providing fly stocks and reagents. Lines obtained from the Bloomington Drosophila Stock Center (NIH P40OD018537) were used in this study. We thank Rui Martinho and Pedro Prudencio for help with experimental protocol optimization. We thank Rui Martinho, Catarina Pereira, Alisson Gontijo, Samuel Walker, Gili Ezra, Ibrahim Tastekin, Dennis Goldschmidt, Patrícia Francisco, Daniel Münch, and all members of the Behavior and Metabolism laboratory for helpful discussions and comments on the manuscript, and Gil Costa for illustrations. We thank Dennis Goldschmidt for typesetting this manuscript, and Ricardo Henriques for the LATEX template. We thank Célia Baltazar, Margarida Anjos and Nicholas Archer for technical assistance. This project was supported by the Portuguese Foundation for Science and Technology (FCT) postdoctoral fellowship SFRH/BPD/79325/2011 to ZCS. Work by ZCS was also financed by national funds through the FCT, in the framework of the financing of the Norma Transitória (DL 57/2016). Research at the Centre for the Unknown is supported by the Champalimaud Foundation.

## Materials & Methods

### *Drosophila* stocks and genetics

Germline expression of transgenes for overexpression or RNAi delivery was achieved by crossing Gal4-carrying female flies (*nanosGal4* (courtesy of Dr. Ralph Neumüller) or *MTD-Gal4* (BL#31777)) with males from the following stocks: UAS*Bam-GFP* (courtesy of Dr. McKearin and Dr. Buszczak), UAS-*Bam*-shRNA (BL#33631), UAS-*HexA*-shRNA (BL#35155), UAS-*Pfk*-shRNA (BL#36782), UAS-*Pyk*shRNA (BL#35218), UAS-*Pgi*-shRNA (BL#51804), UAS*Pgd*-shRNA (BL#65078), UAS-*Zw*-shRNA (BL#50667), UAS-*Rpi*-shRNA (BL#62196), UAS-*GFP*-shRNA (I) (BL#41558), UAS-*GFP*-shRNA (II) (BL#41553) or UAS-*GFP*-shRNA (III) (BL#41552). The RNAi transgene stocks used in this study were originally generated using two different vectors, VALIUM20 or 22, both effective for expression in the germline, and integration in either the attp2 or attp40 site (Perkins et al., 2015). The corresponding control *GFP* knockdown was chosen according to the vector backbone and insertion site of the experimental RNAi line. Double mutant *Pgd* and *Zw* females (*Pgdn*39 *Zwlo*24 (BL#6033)) were crossed to *w*1118 males to generate heterozygous control flies. To test the involvement of *fit* in mediating the anti-satiation effect of the germline a *bam* transgene under the control of a heat shock promoter on the X chromosome (BL#24636) was combined with a *fit* null mutant allele (*Fit*81 courtesy of Dr. Yan Li). To generate homozygous mutant offspring these flies were crossed to the *fit* mutant allele. To generate heterozygous control offspring flies, the same females were crossed to males from the genetic background used to generate the *fit* mutants. The full genotypes of experimental flies are listed in Table S1.

### *Drosophila* rearing, media, and dietary treatments

Flies were reared on yeast-based medium (YBM) (per liter of water: 8 g agar [NZYTech, PT], 80 g barley malt syrup [Próvida, PT], 22 g sugar beet syrup [Grafschafter, DE], 80 g corn flour [Próvida, PT], 10 g soya flour [A. Centazi, PT], 18 g instant yeast [Saf-instant, Lesaffre], 8 ml propionic acid [Argos], and 12 ml nipagin [Tegospet, Dutscher, UK] [15% in 96% ethanol] supplemented with instant yeast granules on the surface [Saf-instant, Lesaffre]). To ensure a homogenous density of offspring among experiments, fly cultures were always set with 6 females and 3 males per vial and left to lay eggs for 3 days. Flies were reared in YBM until adulthood. Holidic media (HM) were prepared as described previously using the FLYaa formulation (Piper et al., 2017), with the exception of the HM used for glucose and trehalose measurements and the two-color food choice assays, for which we used the previous HUNTaa formulation, which only differs in the amount of specific amino acids (Piper et al., 2014). The different HM used in this study are described in Tables S2. Polypropylene fly vials (#734-2261, VWR) were used for rearing the flies in both YBM and HM. In all experiments using the HM, the following dietary treatment protocol was used to ensure a well-fed and mated state: groups of 16 1–5day-old females were collected into fresh YBM-filled vials with 5 *Canton-S* males and transferred to fresh YBM after 48h. Following a period of 24 h, flies were transferred to different HM for 48-72 h and immediately tested in the indicated assay. For the egg laying experiments, groups of 16 1–5-dayold females were collected into fresh YBM-filled vials with 5 *Canton-S* males and transferred to fresh YBM after 48 h. Following a period of 24 h, flies were assayed for egg production. Flies without germline were generated by expressing bam using a heat shock protocol in a water bath at 37°C for 1 h, followed by a 2 h recovery period at 25°C and followed by another heat shock at 37°C for 1 h. This protocol was performed twice at, 6 and 9 days after egg laying. Fly rearing, maintenance, and behavioral testing were performed at 25°C in climate-controlled chambers at 70% relative humidity in a 12-h light–dark cycle (Aralab, FitoClima 60000EH).

### Egg-laying assays

Groups of 16 female and 5 male flies were briefly anesthetized using light CO2 exposure and transferred to apple juice agar plates (per liter, 250 ml apple juice, 19.5 g agar (#MB14801, Nzytech), 20 g sugar, and 10 ml nipagin (10% in ethanol, #25605.293, VWR), where they were allowed to lay eggs for 24 h. Importantly to avoid changes in nutrient state no yeast was added to the egg laying plates. Flies were then removed and counted and the n° eggs assessed. Egg laying was calculated by dividing the number of eggs by the number of living females at the end of the assay.

### Ovary dissection, staining and imaging

Ovaries were dissected in ice cold PBS and fixed with a solution of 4% PFA (#158127, Sigma) and 0.1% Triton X-100 (#21123, Sigma) in PBS for 20 min at RT using soft agitation. Ovaries were washed 3x with 0.1% Triton X-100 in PBS (PBT). Tissue nonspecific antigens were blocked using 0.5% NGS (#16210-064, Invitrogen) dissolved in PBS for 1h at RT and using constant agitation. Ovaries were next incubated with Phalloidin (#P5282, Sigma) at a dilution of 1:25 for 20 min using agitation followed by 3 washes with PBT and 1 wash in PBS before mounting in Vectashield with DAPI (#H-1200, Vector Laboratories). Analysis of the tissue and image acquisition was carried out using a Zeiss LSM 710 confocal laserscanning microscope and processed using Fiji and Adobe Photoshop.

### Generation and preparation of probes for in situ hybridization

RNA probes for *in situ* hybridization were synthesized from a cDNA library derived from wild-type flies (protocol were adapted from Morris et al. (2009). For this, mRNA was extracted from 15 wild type flies (BL#2057) using the following procedure: flies were snap frozen on dry ice before grinded and homogenized for 20 s (using pestles #Z359947, Sigma) in 100 μl of PureZOL (#732-6890, Bio-Rad). 900 μl of PureZOL was further added and mixed by vortexing. 200 μl of chloroform (#C2432, Sigma) were added and the samples were incubated on ice for 15 min. After centrifuging the samples at top speed for 15 min in 4°C (Eppendorf Centrifuge 5415 R), the aqueous phase was transferred to a new RNase free Eppendorf tube. RNA was precipitated by mixing 500 μl of isopropanol (#278475, Sigma) with the aqueous phase. Samples were incubated on ice for 10 min and finally centrifuged at top speed for 10 min at 4°C. The RNA pellet was washed with 500 μl of 75% ethanol and air-dried at RT. The pellet was resuspended in 12 μl of RNase free water. The concentration of the total mRNA samples was determined using a Nanodrop (Thermo Scientific, Nanodrop 2000 Spectophotometer). cDNA was synthesized using the Transcriptor high fidelity cDNA synthesis KIT (#05081955001, Roche) according to manufacturer’s instructions. The DNA of exonic sequences for *HexA* and *Pgd* (retrieved from Ensembl genome browser 93 (https://www.ensembl.org/index.html) for synthesizing the probes was amplified using the primers described in Table S3 as following: the PCR reaction was set with KOD hot start master mix (#71842, Novagen), according to the manufacturer’s instructions and subsequently purified (#28706, Quiagen). In vivo transcription was carried out using the RNA polymerase SP6/T3 (#M0378S/M0207S, NEB) followed by template DNA degradation using DNAseI (#M0303S, NEB) according to manufacturer’s instructions. Probes were hydrolyzed using 20 μl of carbonate buffer (2x) (composition in Table S4) for 20 min at 65°C. 1.67 μl of Lithium Chloride (6M) and 120 μl of ethanol (100%) were added and the RNA probes were left to precipitate O/N at 20°C. Samples were centrifuged for 30 min at top speed in 4°C, and probes were washed in 200 μl of 70% ethanol. The pellets were air-dried and resuspended in 50 μl of hybridization buffer (composition in Table S4).

### Ovary dissection and*in situ* hybridization

Protocol for *in situ* hybridization was adapted from (Morris et al., 2009). Ovaries were dissected in PBS on ice and fixed with a solution of 4% PFA and 0,1% Triton X-100 in PBS for 20 min at RT. Ovaries were washed 3x with 0.3% Tween 20 (#P9416, Sigma) in PBS. Ovaries were next incubated with 200 μl of pre-hybridization buffer (composition in Table S4) for 1 h at RT followed by a 1 h incubation with 200 μl of hybridization buffer at 55°C. Ovaries were left incubating with the probes at 1:100 dilution in hybridization buffer O/N at 55°C. The ovaries were washed in pre-hybridization buffer for 30 min at 55°C, followed by 5 washes with 1% Tween 20 in PBS (PBT) for 20 min each at RT. They were next incubated for 30 min in Roche blocking buffer (1:10 in PBT) (#11096176001, Sigma) at RT followed by incubation with mouse anti-DIG alkaline phosphatase (AP) antibody (1:2000) (#11093274910, Roche) in Roche blockingbuffer (1:10 in PBT) at 4°C. Ovaries were next washed 5 times in PBT at RT, and then rinsed in 1X AP buffer (composition in Table S4). The tissue was stained with NBT/BCIP (1:50) (#11681451001, Roche) in AP buffer using a multiwell plate and checked under the scope regularly. The reaction was stopped after approximately 1 h with PBT, followed by washing in PBT 3 times, and finally mounted in Vectashield (#H1000, Vector Labs). Analysis of the tissue and image acquisition was carried out using Zeiss AxioScan.Z1 automated brightfield slide scanner and processed using Fiji and Adobe Photoshop.

### flyPAD assays

flyPAD assays were performed as described in (Itskov et al., 2014). Single flies maintained in different dietary conditions were tested in arenas that contained two kinds of food patches: 10% yeast and 20 mM sucrose, each mixed with 1% agarose. Flies were individually transferred to flyPAD arenas by mouth aspiration and allowed to feed for 1 h at 25°C, 70% relative humidity. All assays were performed between ZT2 and ZT9. The total number of sips per animal during the assay was calculated using previously described algorithms (Itskov et al., 2014). Flies that did not eat (defined as having fewer than two activity bouts during the assay) were excluded from the analysis.

### Two-color food choice assay

Two-color feeding preference assays were performed as previously described (Ribeiro and Dickson, 2010). Groups of 16 female and 5 male flies were briefly anesthetized using light CO2 exposure and introduced into tight-fit-lid Petri dishes (#351006, Falcon). Flies were given the choice between nine spots of 10 μl sucrose solution mixed with red colorant (20 mM sucrose (#84097, Sigma-Aldrich); 7.5 mg/ml agarose (#16500, Invitrogen); 5 mg/ml Erythrosin B (#198269, Sigma-Aldrich); 10% PBS) and nine spots of 10 μl yeast solution mixed with blue colorant (10% yeast (Saf-instant, Lesaffre); 7.5 mg/ml agarose; 0.25 mg/ml Indigo carmine (#131164, Sigma-Aldrich); 10% PBS) for 2 h. After visual inspection of the abdomen under the stereo microscope (Zeiss, Stereo Discovery.V8), each female fly was scored as having eaten either sucrose (red abdomen), yeast (blue abdomen), or both (red and blue or purple abdomen) media. The sugar preference index (SPI) for the whole female population in the assay was calculated a follows: (*n*red sucrose *− n*blue yeast) / (*n*red sucrose + *n*blue yeast + *n*both). Dye-swap (red yeast versus blue sucrose choice) experiments has been tested previously (Leitao-Goncalves et al., 2017) and because the colorant used for each food source had no impact on overall feeding preference, we opted to exclusively perform red sucrose versus blue yeast choice experiments. All assays were performed between ZT6 and ZT9. In all experiments, the observer was blind for both diet and genotype.

### Carbohydrate measurements

The glucose/trehalose measurement protocol was adapted from (Tennessen et al., 2014). Concentrations of these metabolites were measured from 15 fly heads and in 5-6 replicates per condition. Flies were harvested and washed in PBS before snap frozen in liquid nitrogen. Fly heads were separated from bodies using sieves of appropriate sizes (#11342204, Fisher Scientific). The tissue was grinded and homogenized in 100 μl of cold Trehalase buffer (TB) (composition in Table S4) (using pestles, #Z359947, Sigma). 15 μl of homogenate was used for protein measurement using a Pierce BCA protein Assay Kit (#10678484, Fisher Scientific) and a standard curve was calculated with a series of Albumin dilutions (0.025, 0.125, 0.250, 0.5, 0.75, 1, 1.5 and 2 mg/ml). Absorbance was measured at 562 nm in a plate reader (BMG Labtech, SpectroStar Nano) and the concentrations were calculated from the albumin standard curve. The remaining homogenate was heated for 10 min at 70°C and spun for 3 min at top speed at 4°C (Eppendorf 5415R). The supernatant was transferred to a new tube and heated for 10 min at 70°C. Samples were then centrifuged at top speed at 4°C. The supernatant was transferred to clean tubes. To generate standard curves, glucose (#GAHK-20, Sigma) dilution series were prepared in TB (0.16, 0.08, 0.04, 0.02 and 0.01 mg/ml) and trehalose (#T0167, Sigma) dilution series in a 1:1 mix of TB and Trehalase (3μl trehalase/ml, TS) (#T8778, Sigma) (0.16, 0.08, 0.04, 0.02 and 0.01 mg/ml). 30 μl of each sample was mixed either with 30 μl of TB (used to calculate the free glucose) or 30 μl of TS (used to calculate the trehalose derived glucose). 30 μl of the glucose standards were mixed with 30 μl of TB, and 30 μl of trehalose standards were mixed either with 30 μl of TB or 30 μl of TS. All standards and samples were incubated at 37°C for 18-24h. 30 μl of each sample and standards were transferred to a 96-well plate, mixed with 100 μl of the Glucose assay reagent (#GAHK-20, Sigma) and incubated for 1 h at RT. Absorbance was measured at 340 nm in a plate reader (BMG Labtech, SpectroStar Nano). A blank sample was prepared for glucose measurements using ddH_2_O and TB and a blank sample for the trehalose measurements was prepared with ddH_2_O and TS. Blank sample absorbances were subtracted from the glucose or the trehalose samples. Free glucose concentrations were calculated using the absorbances from the undigested samples and using the glucose standard curve. The trehalose concentrations were calculated by subtracting the absorbance of the undigested samples from the digested samples, and then using the trehalose standard curve. Concentrations of these metabolites in each sample were normalized for the corresponding protein content. Fructose measurement protocol was adapted from (Miyamoto et al., 2012). Measurements were performed from 60 fly heads and in 3 replicates per condition. Flies were harvested and washed in PBS before being snap frozen in liquid nitrogen. Fly heads were separated from bodies using sieves of appropriate sizes (#11342204, Fisher Scientific). Fly heads were grinded and homogenized in 100 μl of cold ddH_2_O (using pestles #Z359947, Sigma) and ddH_2_0 was further added to obtain a total volume of 175 μl. 15 μl of homogenate was used for calculating protein concentration as described in the glucose/trehalose measurement protocol. To generate a standard curve, fructose dilutions were prepared in ddH_2_O (400, 200, 100, 50, 25, 12.5 μg/ml) (#F0127, Sigma). 2 times the sample volume of 0.05% Resorcinol (#10149831, Fisher Scientific) in HCl (6N) (#30721, MERCK) were added to each sample and fructose dilutions. Samples were centrifuged at top speed for 10 min and 200 μl of the supernatant was transferred to 2 clean tubes. One of the tubes was heated for 10 min at 95°C, the other tube was used for background absorbance measurement. Two blank samples were prepared using ddH_2_O and Resorcinol, one of which was heated. Absorbance was measured at 485 nm using a plate reader (BMG Labtech, SpectroStar Nano). The average absorbance of the non-heated blank was subtracted from the non-heated samples and the same procedure was done for the heated samples. Finally, the absorbance measurements of the non-heated samples and standard dilutions were subtracted from the corresponding heated samples. A standard curve was calculated from the fructose dilutions and the concentrations of fructose in each sample was calculated using this curve. Protein measurements were performed as described in the previous section and the fructose concentrations normalized to protein amounts.

### Starvation assay

Groups of 16 1–5-day-old females were collected into fresh YBM-filled vials with 5 *Canton-S* males and transferred to fresh YBM after 48 h to ensure they were well fed and mated. After one day, single flies were transferred to tubes containing water soaked paper for a total of 80 flies per condition. The number of living flies was scored every 12 h until all flies were dead.

### PER assay

PER assays were performed as described in (Walker et al., 2015). Briefly, groups of 16 1–5-day-old females were collected into fresh YBM-filled vials with 5 *Canton-S* males and transferred to fresh YBM after 48 h to ensure they were well fed and mated. After one day, flies were then gently anaesthetized using CO2 and affixed by the dorsal thorax to a glass slide using No More Nails (UniBond) in groups of 20 for a total of 35-40 flies tested. Flies were allowed to recover for 2 hr at 25°C in a humidified box and then moved to room temperature. They were first allowed to drink water until they no longer responded to stimulation, and then a droplet of 25 mM sucrose (#84097, Sigma) was presented for 3 s on the tarsi. Flies were scored as extending versus not extending the proboscis and each fly was treated as a single data point for each stimulus.

### Total mRNA extraction, RT-PCR, and quantitative real-time PCR

Flies used for mRNA extraction were snap frozen in dry ice and strored at –80°C. Behavioral assays were performed in parallel to confirm that sibling flies presented the expected feeding phenotype. mRNA was extracted from flies (5 flies per condition) using the following procedure: flies were grinded and homogenized for 20 s (using pestles #Z359947, Sigma) in 100 μl of PureZOL (#732-6890, BioRad). 900 μl of PureZOL was further added and mixed using a vortexer. Subsequently, 200 μl of chloroform was added to the samples, followed by 15 s vigorous shaking, incubation on ice for 15 min, and finally centrifuged at top speed for 15 min at 4°C. The aqueous phase was transferred to a new tube and the RNA was precipitated by mixing with 500 μl of isopropanol (#278475, Sigma). The samples were stored on ice for 10 min and spun at top speed for 10 min at 4°C. The supernatant was removed and the RNA pellet washed with 500 μl of 75% ethanol. After one last centrifugation at top speed for 10 min at 4°C, the ethanol was removed and the pellet was air-dried. RNA was resuspended in 12 μl of distilled RNase/DNase-free water. After RNA quantification using a Nanodrop (Thermo Scientific, Nanodrop 2000 Spectophotometer), 1 μg of RNA was reverse transcribed (RT) using the iScript Reverse Transcription Supermix for RT-PCR kit (#170-8840 Bio-Rad), following the manufacturer’s instructions. The expression of *fit* was determined using realtime PCR. Each cDNA sample was amplified using SsoFast EvaGreen Supermix on the CFX96 Real-Time System (BioRad). Briefly, the reaction conditions consisted of 1 μl of cDNA, 1 μl (10 μM) of each primer, 10 μl of supermix, and 7 μl of water. The cycle program consisted of enzyme activation at 95°C for 30 s, 39 cycles of denaturation at 95°C for 2 s, and annealing and extension for 5 s. The primers used in this reaction are listed in Table S3. Three experimental replicas and two technical replicas per genotype were used. Appropriate non-template controls were included in each 96well PCR reaction, and dissociation analysis was performed at the end of each run to confirm the specificity of the reaction. Absolute levels of RNA were calculated from a standard curve and normalized to two internal controls (*Actin42A* and *RpL32*). The relative quantitation of each mRNA was performed using the comparative Ct method. Data processing was performed using Bio-rad CFX Manager 3.1 (Bio-Rad).

## Bibliography

Andermann ML, Lowell BB, 2017. Toward a Wiring Diagram Understanding of Appetite Control. Neuron 95(4):757–778. doi:10.1016/j.neuron.2017.06.014

Barton LJ, LeBlanc MG, Lehmann R, 2016. Finding their way: themes in germ cell migration. Current Opinion in Cell Biology 42:128–137. doi:10.1016/j.ceb.2016.07.007

Bastock R, Johnston DS, 2008. Drosophila oogenesis. Current Biology 18(23):R1082–R1087. doi:10.1016/j.cub.2008.09.011

Bryant M, Truesdale KP, Dye L, 2006. Modest changes in dietary intake across the menstrual cycle: implications for food intake research. British Journal of Nutrition 96(5):888–894. doi:10.1017/BJN20061931

Carvalho-Santos Z, Ribeiro C, 2018. Gonadal ecdysone titers are modulated by protein availability but do not impact protein appetite. Journal of Insect Physiology 106(Pt 1):30–35. doi:10.1016/j.jinsphys.2017.08.006

Cavener DR, 1980. Genetics of male-specific glucose oxidase and the identification of other unusual hexose enzymes in Drosophila melanogaster. Biochemical genetics 18(9–10):929–37

Chintapalli VR, Wang J, Dow JA, 2007. Using FlyAtlas to identify better Drosophila melanogaster models of human disease. Nature genetics 39(6):715–20. doi:10.1038/ng2049

Coll AP, Farooqi IS, O’Rahilly S, 2007. The Hormonal Control of Food Intake. Cell 129(2):251–262. doi:10.1016/j.cell.2007.04.001

Colombani J, Andersen DS, Leopold P, 2012. Secreted Peptide Dilp8 Coordinates Drosophila Tissue Growth with Developmental Timing. Science 336(6081):582–585. doi:10.1126/science.1216689

Corrales-Carvajal VM, Faisal AA, Ribeiro C, 2016. Internal states drive nutrient homeostasis by modulating exploration-exploitation trade-off. Elife 5. doi:10.7554/eLife.19920

Cox RT, Spradling AC, 2003. A Balbiani body and the fusome mediate mitochondrial inheritance during Drosophila oogenesis. Development 130(8):1579–1590. doi:10.1242/dev.00365

de Cuevas M, Lilly M, Spradling A, 1997. Germline Cyst Formation in Drosophila. Annual Review of Genetics 31(1):405–428. doi:10.1146/annurev.genet.31.1.405

DeBerardinis RJ, Chandel NS, 2016. Fundamentals of cancer metabolism. Science Advances 2(5):e1600200. doi:10.1126/sciadv.1600200

DeBerardinis RJ, Lum JJ, Hatzivassiliou G, Thompson CB, 2008a. The Biology of Cancer: Metabolic Reprogramming Fuels Cell Growth and Proliferation. Cell Metabolism 7(1):11–20. doi:10.1016/j.cmet.2007.10.002

DeBerardinis RJ, Sayed N, Ditsworth D, Thompson CB, 2008b. Brick by brick: metabolism and tumor cell growth. Current Opinion in Genetics & Development 18(1):54–61. doi:10.1016/j.gde.2008.02.003

Dethier VG, 1976. The hungry fly: A physiological study of the behavior associated with feeding. Harvard U Press, Oxford, England

Djabrayan NJV, Smits CM, Krajnc M, Stern T, Yamada S, Lemon WC, Keller PJ, Rushlow CA, Shvartsman SY, 2019. Metabolic Regulation of Developmental Cell Cycles and Zygotic Transcription. Current Biology 29(7):1193–1198.e5. doi:10.1016/j.cub.2019.02.028

Droujinine IA, Perrimon N, 2016. Interorgan Communication Pathways in Physiology: Focus on Drosophila. Annual review of genetics 50:539–570. doi:10.1146/annurev-genet-121415-122024

Drummond-Barbosa D, Spradling AC, 2001. Stem cells and their progeny respond to nutritional changes during Drosophila oogenesis. Dev Biol 231(1):265–78. doi:10.1006/dbio.2000.0135

Dumollard R, Duchen M, Carroll J, 2007. The Role of Mitochondrial Function in the Oocyte and Embryo. In Current Topics in Developmental Biology, volume 77 of The Mitochondrion in the Germline and Early Development, pp. 21–49. Academic Press. DOI:10.1016/S0070-2153(06)77002-8

Dye L, Blundell JE, 1997. Menstrual cycle and appetite control: implications for weight regulation. Human Reproduction 12(6):1142–1151. doi:10.1093/humrep/12.6.1142

Eisenreich W, Ettenhuber C, Laupitz R, Theus C, Bacher A, 2004. Isotopolog perturbation techniques for metabolic networks: Metabolic recycling of nutritional glucose in Drosophila melanogaster. Proceedings of the National Academy of Sciences 101(17):6764–6769. doi:10.1073/pnas.0400916101

Friedman JM, Halaas JL, 1998. Leptin and the regulation of body weight in mammals. Nature 395(6704):763–770. doi:10.1038/27376

Fujii S, Amrein H, 2002. Genes expressed in the Drosophila head reveal a role for fat cells in sex-specific physiology. The EMBO journal 21(20):5353–63

Fujikawa K, Takahashi A, Nishimura A, Itoh M, Takano-Shimizu T, Ozaki M, 2009. Characteristics of genes up-regulated and down-regulated after 24 h starvation in the head of Drosophila. Gene 446(1):11–7. doi:10.1016/j.gene.2009.06.017

Gambhir SS, 2002. Molecular imaging of cancer with positron emission tomography. Nature Reviews Cancer 2(9):683. doi:10.1038/nrc882

Garelli A, Gontijo AM, Miguela V, Caparros E, Dominguez M, 2012. Imaginal Discs Secrete Insulin-Like Peptide 8 to Mediate Plasticity of Growth and Maturation. Science 336(6081):579–582. doi:10.1126/science.1216735

Giese GE, Nanda S, Holdorf AD, Walhout AJM, 2019. Transcriptional regulation of metabolic flux: A Caenorhabditis elegans perspective. Current Opinion in Systems Biology 15:12–18. doi:10.1016/j.coisb.2019.03.002

Goncalves MD, Hopkins BD, Cantley LC, 2019a. Dietary Fat and Sugar in Promoting Cancer Development and Progression. Annual Review of Cancer Biology 3(1):255–273. doi:10.1146/annurev-cancerbio-030518-055855

Goncalves MD, Lu C, Tutnauer J, Hartman TE, Hwang SK, Murphy CJ, Pauli C, Morris R, Taylor S, Bosch K, Yang S, Wang Y, Van Riper J, Lekaye HC, Roper J, Kim Y, Chen Q, Gross SS, Rhee KY, Cantley LC, Yun J, 2019b. High-fructose corn syrup enhances intestinal tumor growth in mice. Science 363(6433):1345–1349. doi:10.1126/science.aat8515

Gvozdev VA, Gerasimova TI, Kogan GL, Braslavskaya O, 1976. Role of the pentose phosphate pathway in metabolism of Drosophila melanogaster elucidated by mutations affecting glucose 6-phosphate and 6-phosphogluconate dehydrogenases. FEBS letters 64(1):85–8

Heiden MGV, DeBerardinis RJ, 2017. Understanding the Intersections between Metabolism and Cancer Biology. Cell 168(4):657–669. doi:10.1016/j.cell.2016.12.039

Hirabayashi S, Baranski T, Cagan R, 2013. Transformed Drosophila Cells Evade Diet-Mediated Insulin Resistance through Wingless Signaling. Cell 154(3):664–675. doi:10.1016/j.cell.2013.06.030

Hsu HJ, Drummond-Barbosa D, 2009. Insulin levels control female germline stem cell maintenance via the niche in Drosophila. Proc Natl Acad Sci U S A 106(4):1117–21. doi:10.1073/pnas.0809144106

Hughes MB, Lucchesi JC, 1977. Genetic rescue of a lethal “null” activity allele of 6-phosphogluconate dehydrogenase in Drosophila melanogaster. Science 196(4294):1114–5

Inagaki HK, de Leon SBT, Wong AM, Jagadish S, Ishimoto H, Barnea G, Kitamoto T, Axel R, Anderson DJ, 2012. Visualizing Neuromodulation In Vivo: TANGO-Mapping of Dopamine Signaling Reveals Appetite Control of Sugar Sensing. Cell 148(3):583–595. doi:https://doi.org/10.1016/j.cell.2011.12.022

Itskov PM, Moreira JM, Vinnik E, Lopes G, Safarik S, Dickinson MH, Ribeiro C, 2014. Automated monitoring and quantitative analysis of feeding behaviour in Drosophila. Nat Commun 5:4560. doi:10.1038/ncomms5560

Itskov PM, Ribeiro C, 2013. The Dilemmas of the Gourmet Fly: The Molecular and Neuronal Mechanisms of Feeding and Nutrient Decision Making in Drosophila. Frontiers in Neuroscience 7. doi:10.3389/fnins.2013.00012

Johnston DS, Ahringer J, 2010. Cell Polarity in Eggs and Epithelia: Parallels and Diversity. Cell 141(5):757–774. doi:10.1016/j.cell.2010.05.011

Lehmann R, 2012. Germline Stem Cells: Origin and Destiny. Cell Stem Cell 10(6):729–739. doi:10.1016/j.stem.2012.05.016

Leitao-Goncalves R, Carvalho-Santos Z, Francisco AP, Fioreze GT, Anjos M, Baltazar C, Elias AP, Itskov PM, Piper MDW, Ribeiro C, 2017. Commensal bacteria and essential amino acids control food choice behavior and reproduction. PLoS biology 15(4):e2000862. doi:10.1371/journal.pbio.2000862

Leopold P, Perrimon N, 2007. Drosophila and the genetics of the internal milieu. Nature 450(7167):186–188. doi:10.1038/nature06286

Liu B, Winkler F, Herde M, Witte CP, Großhans J, 2019. A Link between Deoxyribonucleotide Metabolites and Embryonic Cell-Cycle Control. Current Biology 29(7):1187–1192.e3. doi:10.1016/j.cub.2019.02.021

Lunt SY, Vander Heiden MG, 2011. Aerobic glycolysis: meeting the metabolic requirements of cell proliferation. Annu Rev Cell Dev Biol 27:441–64. doi:10.1146/annurev-cellbio-092910-154237

Malmanche N, Clark DV, 2004. Drosophila melanogaster Prat, a Purine de Novo Synthesis Gene, Has a Pleiotropic Maternal-Effect Phenotype. Genetics 168(4):2011–2023. doi:10.1534/genetics.104.033134

McLaughlin JM, Bratu DP, 2015. Drosophila melanogaster Oogenesis: An Overview. In Bratu DP, McNeil GP, eds., Drosophila Oogenesis: Methods and Protocols, Methods in Molecular Biology, pp. 1–20. Springer New York, New York, NY. DOI:10.1007/978-1-4939-28514_1

Min KJ, Hogan MF, Tatar M, O’Brien DM, 2006. Resource allocation to reproduction and soma in Drosophila: a stable isotope analysis of carbon from dietary sugar. Journal of Insect Physiology 52(7):763–70. doi:10.1016/j.jinsphys.2006.04.004

Miyamoto T, Slone J, Song X, Amrein H, 2012. A fructose receptor functions as a nutrient sensor in the Drosophila brain. Cell 151(5):1113–1125. doi:10.1016/j.cell.2012.10.024

Morris CA, Benson E, White-Cooper H, 2009. Determination of gene expression patterns using in situ hybridization to Drosophila testes. Nature Protocols 4(12):1807–1819. doi:10.1038/nprot.2009.192

O’Brien DM, Min KJ, Larsen T, Tatar M, 2008. Use of stable isotopes to examine how dietary restriction extends Drosophila lifespan. Current biology: CB 18(4):R155–6. doi:10.1016/j.cub.2008.01.021

Ohlstein B, McKearin D, 1997. Ectopic expression of the Drosophila Bam protein eliminates oogenic germline stem cells. Development 124(18):3651–62

Parisi MJ, Gupta V, Sturgill D, Warren JT, Jallon JM, Malone JH, Zhang Y, Gilbert LI, Oliver B, 2010. Germline-dependent gene expression in distant non-gonadal somatic tissues of Drosophila. BMC genomics 11:346. doi:10.1186/1471-2164-11-346

Pavlova NN, Thompson CB, 2016. The Emerging Hallmarks of Cancer Metabolism. Cell metabolism 23(1):27–47. doi:10.1016/j.cmet.2015.12.006

Perkins LA, Holderbaum L, Tao R, Hu Y, Sopko R, McCall K, Yang-Zhou D, Flockhart I, Binari R, Shim HS, Miller A, Housden A, Foos M, Randkelv S, Kelley C, Namgyal P, Villalta C, Liu LP, Jiang X, Huan-Huan Q, Wang X, Fujiyama A, Toyoda A, Ayers K, Blum A, Czech B, Neumuller R, Yan D, Cavallaro A, Hibbard K, Hall D, Cooley L, Hannon GJ, Lehmann R, Parks A, Mohr SE, Ueda R, Kondo S, Ni JQ, Perrimon N, 2015. The Transgenic RNAi Project at Harvard Medical School: Resources and Validation. Genetics 201(3):843–852. doi:10.1534/genetics.115.180208

Piper MD, Blanc E, Leitao-Goncalves R, Yang M, He X, Linford NJ, Hoddinott MP, Hopfen C, Soultoukis GA, Niemeyer C, Kerr F, Pletcher SD, Ribeiro C, Partridge L, 2014. A holidic medium for Drosophila melanogaster. Nat Methods 11(1):100–5. doi:10.1038/nmeth.2731

Piper MD, Soultoukis GA, Blanc E, Mesaros A, Herbert SL, Juricic P, He X, Atanassov I, Salmonowicz H, Yang M, Simpson SJ, Ribeiro C, Partridge L, 2017. Matching Dietary Amino Acid Balance to the In Silico-Translated Exome Optimizes Growth and Reproduction without Cost to Lifespan. Cell metabolism 25(3):610–621. doi:10.1016/j.cmet.2017.02.005

Pool LJ, Scott K, 2014. Feeding regulation in Drosophila. Current opinion in neurobiology 29:57–63. doi:10.1016/j.conb.2014.05.008

Ribeiro C, Dickson BJ, 2010. Sex peptide receptor and neuronal TOR/S6K signaling modulate nutrient balancing in Drosophila. Current biology: CB 20(11):1000–5. doi:10.1016/j.cub.2010.03.061

Shyh-Chang N, Daley GQ, Cantley LC, 2013. Stem cell metabolism in tissue development and aging. Development 140(12):2535–47. doi:10.1242/dev.091777

Sieber MH, Spradling AC, 2015. Steroid Signaling Establishes a Female Metabolic State and Regulates SREBP to Control Oocyte Lipid Accumulation. Current Biology 25(8):993–1004. doi:10.1016/j.cub.2015.02.019

Sieber MH, Spradling AC, 2017. The role of metabolic states in development and disease. Curr Opin Genet Dev 45:58–68. doi:10.1016/j.gde.2017.03.002

Sieber MH, Thomsen MB, Spradling AC, 2016. Electron Transport Chain Remodeling by GSK3 during Oogenesis Connects Nutrient State to Reproduction. Cell 164(3):420–432. doi:10.1016/j.cell.2015.12.020

Simpson SJ, Couteur DGL, Raubenheimer D, 2015. Putting the Balance Back in Diet. Cell 161(1):18–23. doi:10.1016/j.cell.2015.02.033

Simpson SJ, Raubenheimer D, 2012. The nature of nutrition. Princeton, Princeton University Press

Simpson SJ, Sword GA, Lorch PD, Couzin ID, 2006. Cannibal crickets on a forced march for protein and salt. Proceedings of the National Academy of Sciences 103(11):4152–4156. doi:10.1073/pnas.0508915103

Slaidina M, Lehmann R, 2014. Translational control in germline stem cell development. The Journal of Cell Biology 207(1):13–21. doi:10.1083/jcb.201407102

Søndergaard L, Mauchline D, Egetoft P, White N, Wulff P, Bownes M, 1995. Nutritional response in a Drosophila yolk protein gene promoter. Molecular and General Genetics MGG 248(1):25–32. doi:10.1007/BF02456610

Song Y, Marmion RA, Park JO, Biswas D, Rabinowitz JD, Shvartsman SY, 2017. Dynamic Control of dNTP Synthesis in Early Embryos. Developmental Cell 42(3):301–308.e3. doi:10.1016/j.devcel.2017.06.013

Steck K, Walker SJ, Itskov PM, Baltazar C, Moreira JM, Ribeiro C, 2018. Internal amino acid state modulates yeast taste neurons to support protein homeostasis in Drosophila. Elife 7. doi:10.7554/eLife.31625

Stincone A, Prigione A, Cramer T, Wamelink MMC, Campbell K, Cheung E, Olin-Sandoval V, Grüning NM, Krüger A, Tauqeer Alam M, Keller MA, Breitenbach M, Brindle KM, Rabinowitz JD, Ralser M, 2015. The return of metabolism: biochemistry and physiology of the pentose phosphate pathway. Biological Reviews 90(3):927–963. doi:10.1111/brv.12140

Stryer L, 1995. Biochemistry. W. H. Freeman and Company, New York, 4th edition

Sun J, Liu C, Bai X, Li X, Li J, Zhang Z, Zhang Y, Guo J, Li Y, 2017. Drosophila FIT is a protein-specific satiety hormone essential for feeding control. Nat Commun 8:14161. doi:10.1038/ncomms14161

Teixeira FK, Sanchez CG, Hurd TR, Seifert JRK, Czech B, Preall JB, Hannon GJ, Lehmann R, 2015. ATP synthase promotes germ cell differentiation independent of oxidative phosphorylation. Nature Cell Biology 17(5):689–696. doi:10.1038/ncb3165

Tennessen JM, Barry WE, Cox J, Thummel CS, 2014. Methods for studying metabolism in Drosophila. Methods 68(1):105–115. doi:10.1016/j.ymeth.2014.02.034

Than TT, Delay ER, Maier ME, 1994. Sucrose threshold variation during the menstrual cycle. Physiology & Behavior 56(2):237–239. doi:10.1016/0031-9384(94)90189-9

Trumper S, Simpson SJ, 1993. Regulation of salt intake by nymphs of Locusta migratoria. Journal of Insect Physiology 39(10):857–864. doi:10.1016/0022-1910(93)90118-B

Vander Heiden MG, Cantley LC, Thompson CB, 2009. Understanding the Warburg effect: the metabolic requirements of cell proliferation. Science 324(5930):1029–33. doi:10.1126/science.1160809

Walker SJ, Corrales-Carvajal VM, Ribeiro C, 2015. Postmating Circuitry Modulates Salt Taste Processing to Increase Reproductive Output in Drosophila. Current biology: CB 25(20):2621–30. doi:10.1016/j.cub.2015.08.043

Walker SJ, Goldschmidt D, Ribeiro C, 2017. Craving for the future: the brain as a nutritional prediction system. Current Opinion in Insect Science 23(Supplement C):96–103. doi:10.1016/j.cois.2017.07.013

Wallace RA, Selman K, 1990. Ultrastructural aspects of oogenesis and oocyte growth in fish and amphibians. Journal of Electron Microscopy Technique 16(3):175–201. doi:10.1002/jemt.1060160302

Warburg O, 1956. On the Origin of Cancer Cells. Science 123(3191):309–314. doi:10.1126/science.123.3191.309

Warburg O, Christian W, 1936. Optischer Nachweis der Hydrierung und Dehydrierung des Pyridins im Gärungs-Co-Ferment. Biochemische Zeitschrift 286:81–81

Warburg O, Christian W, Griese A, 1935. Wasserstoffübertragendes CoFerment, seine Zusammensetzung und Wirkungsweise. Biochemische Zeitschrift 282:157–205

Warburg O, Wind F, Negelein E, 1926. Über den Stoffwechsel von Tumoren im Körper. Klinische Wochenschrift 5(19):829–832. doi:10.1007/BF01726240

Webb P, 1986. 24-hour energy expenditure and the menstrual cycle. The American Journal of Clinical Nutrition 44(5):614–619. doi:10.1093/ajcn/44.5.614

Williams KW, Elmquist JK, 2012. From neuroanatomy to behavior: central integration of peripheral signals regulating feeding behavior. Nature neuroscience 15(10):1350–5. doi:10.1038/nn.3217

