## Supplemental data Santos et a for "Cellular metabolic reprogramming controls sugar appetite"

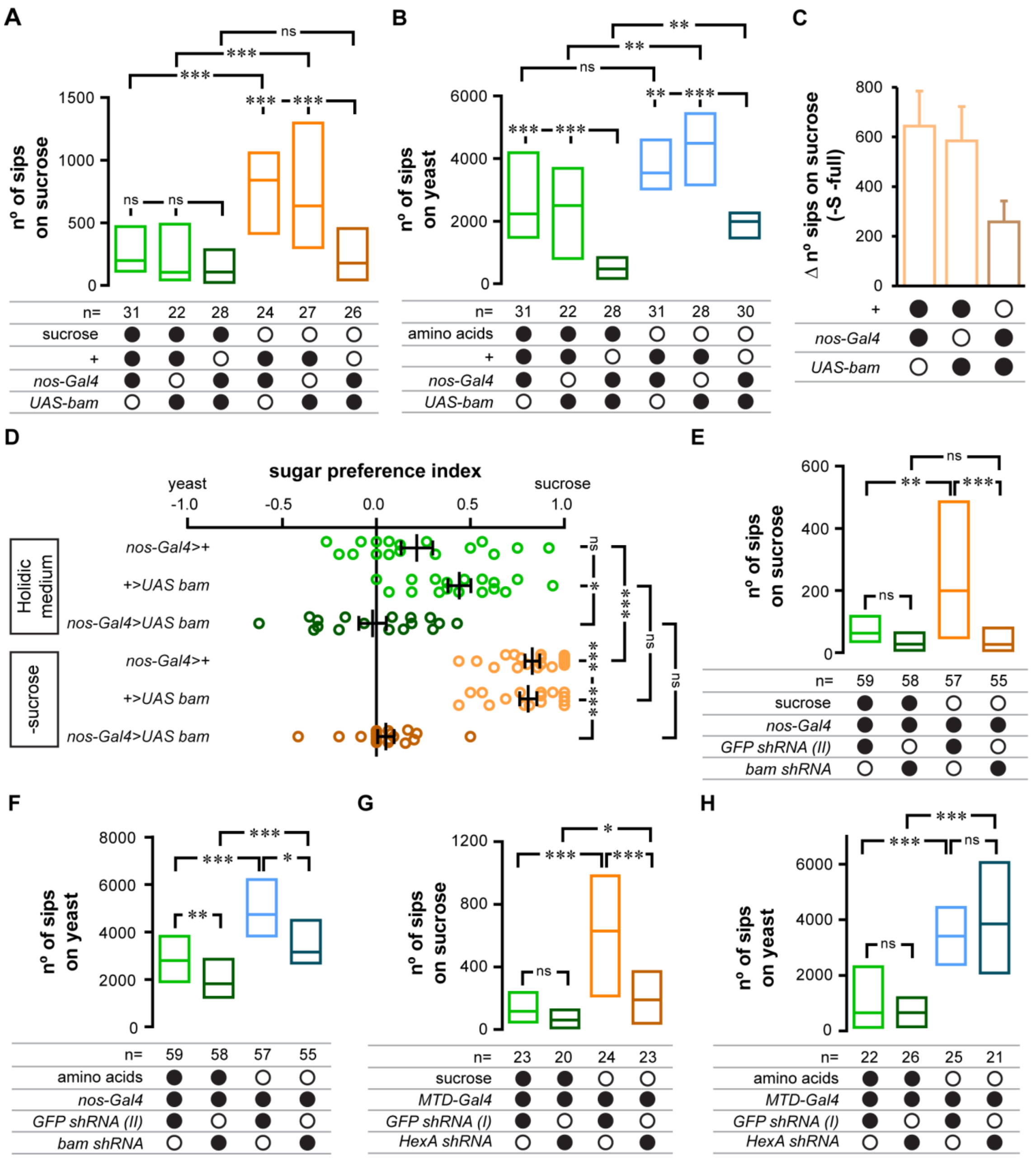


**Fig. S1. Related to Fig. 3. Germline-ablated or females in which germline was metabolically manipulated show a specific decrease in sugar appetite.** (A, B, E-H) Flies were assayed for feeding behavior using the flyPAD technology after being fed for 2 days on full holidic medium (green) or one lacking sucrose (orange) (A, E and G) or amino acids (blue) (B, F and H). *nos-Gal4* or *MTD-Gal4* drivers were used to drive transgene or short hairpin RNAs expression in the germline. *GFP* knockdown lines were used as controls in (E-H). n = number of flies assayed per condition. (C) Virgin females were assayed for an effect in nutrient specific feeding after 2 days on either a complete holidic medium or one lacking sucrose. Sucrose appetite is represented as the difference in sucrose feeding of flies maintained on holidic medium lacking sucrose *vs* full holidic medium. The error bars show 95% confidence interval. (A-C, E-H) Boxes represent median with upper/lower quartiles. Black filled circles represent the presence and open black circles represent the absence of a particular transgene or nutrient in the diet. (D) Sugar preference of flies assayed using the red blue food choice assay kept on full holidic medium (green) or medium lacking sucrose for 2 days (orange). Colored circles in the plot represent sugar preference in single assays, with a line representing the median and error bars representing the interquartile range. n = 17-19. Full genotypes of the flies used in these experiments can be found in **Table S1**. (A, B, D-H) Statistical significance was tested using the Kruskal-Wallis test followed by Dunn's multiple comparison test. ns $p \geq 0.05$, * $p < 0.05$, ** $p < 0.01$, *** $p < 0.001$.

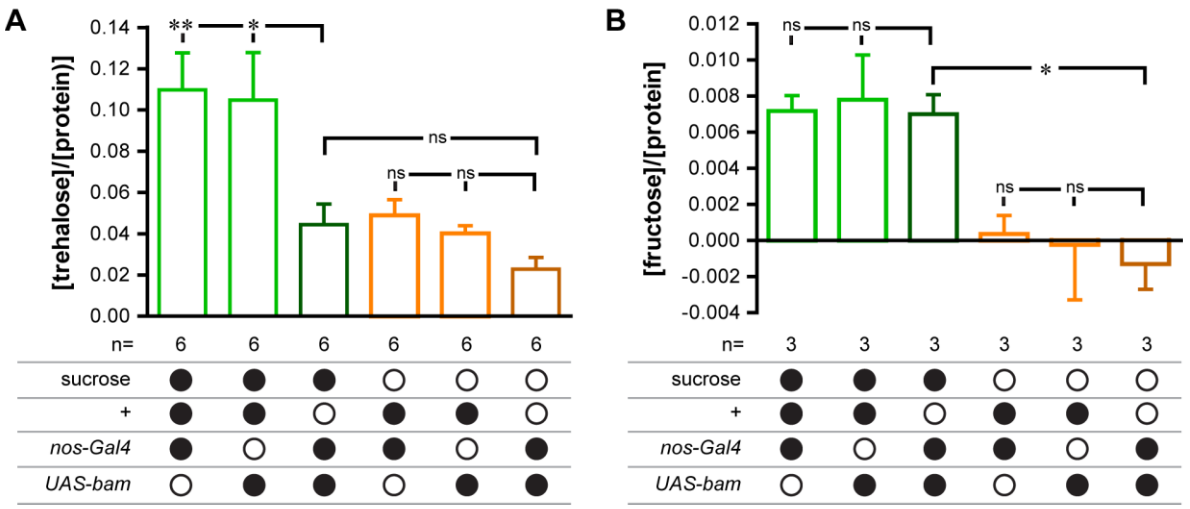


**Fig. S2. Related to Fig. 4. Germline-ablated females do not have increased levels of trehalose or fructose.** Trehalose (A) or fructose (B) measurements from the heads of females fed on full holidic medium (green) or medium lacking sucrose (orange) for 2 days. Sugar concentrations were normalized to protein concentrations in the sample. The columns represent the mean and the error bars the standard error of the mean. n = number of samples used per condition. Statistical significance was tested using an ordinary one-way ANOVA followed by Sidak's multiple comparisons test. ns $p \geq 0.05$, * $p < 0.05$, ** $p < 0.01$.

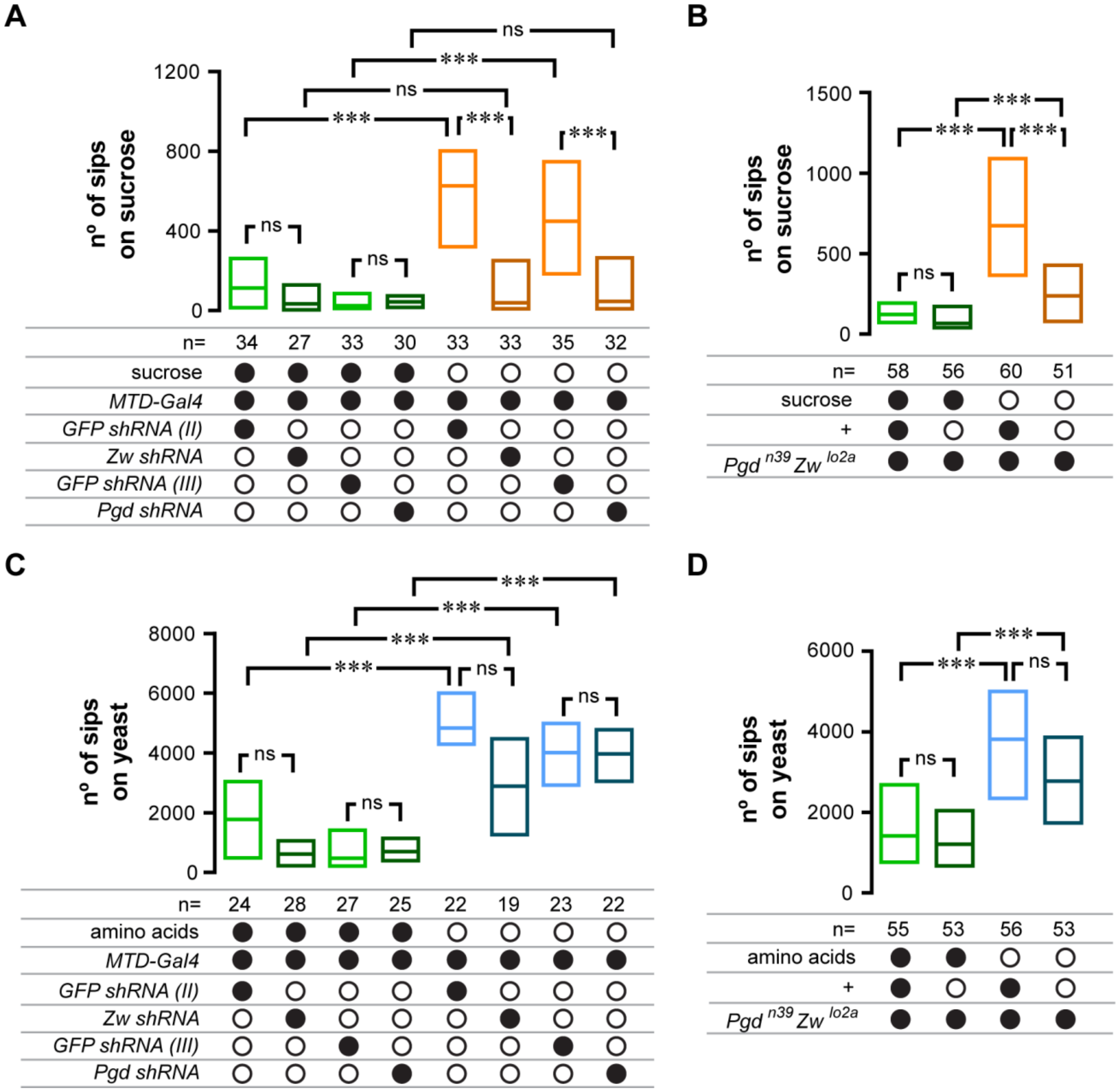


**Fig. S3. Related to Fig. 6. PPP activity in the germline modulates sugar appetite.** Females were assayed for an effect in nutrient specific feeding using the flyPAD technology after 2 days on either a complete holidic medium (green), or one lacking sucrose (orange) (A and B) or amino acids (C and D). Sucrose appetite is represented as the difference in sucrose feeding of flies maintained on holidic medium lacking sucrose *vs* full holidic medium. *MTD-Gal4* driver was used to drive short hairpin RNAs expression in the germline. *GFP* knockdown lines were used as controls. Boxes represent median with upper/lower quartiles. Black filled circles represent the presence and open black circles represent the absence of a particular transgene or nutrient in the diet. n = number of flies assayed per condition. Full genotypes of the flies used in these experiments can be found in **Table S1**. Statistical significance was tested using the Kruskal-Wallis test followed by Dunn's multiple comparison test. ns $p \geq 0.05$, * $p < 0.05$, ** $p < 0.01$, *** $p < 0.001$.

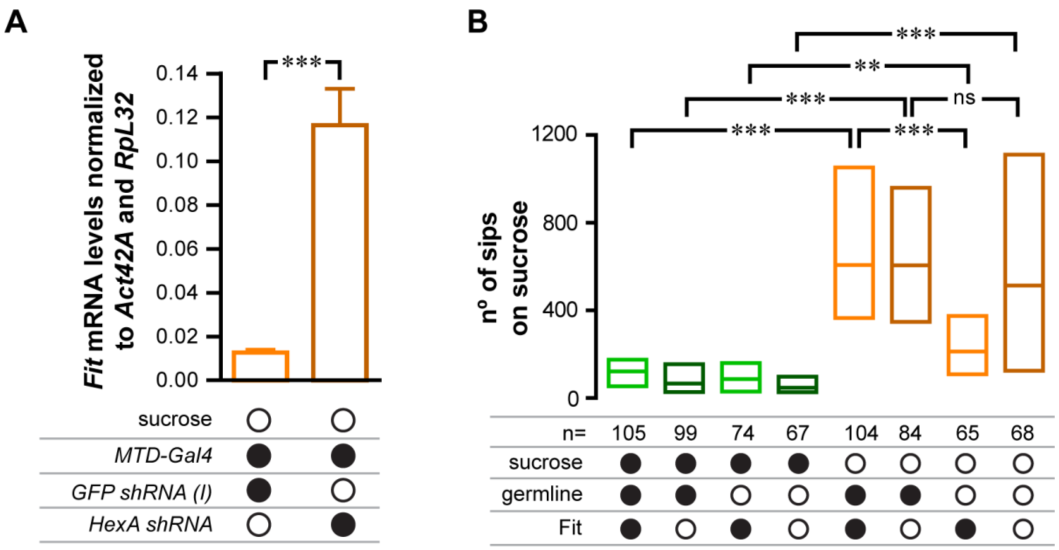


**Fig. S4. Related to Fig. 7. The PPP in the germline controls the expression levels of *Fit*.** (A) *fit* mRNA levels were measured from whole flies fed on holidic medium lacking sucrose and normalized to two internal controls (*Actin 42A* and *RpL32*). *MTD-Gal4* driver was used to drive short hairpin RNA expression for *HexA* in the germline. A genotype matched *GFP* knockdown line was used as a control. (B) Mated female flies were assayed for an effect in nutrient specific feeding using the flyPAD technology after 2 days on either a complete holidic medium (green), or one lacking sucrose (orange). Black filled circles represent the presence and open black circles represent the absence of a transgene, nutrient, the germline or the *fit* gene. n = total number of flies assayed per condition. Statistical significance was tested using the unpaired t test (A) or using the Kruskal-Wallis test followed by Dunn's multiple comparison test (B). In (A) the columns represent the mean and the error bars the standard error of the mean. n = 6. In (B) boxes represent median with upper/lower quartiles Full genotypes of the flies used in these experiments can be found in **Table S1**. ns $p \geq 0.05$, ** $p < 0.01$, *** $p < 0.001$.

**Table S1** – Complete genotypes of the fly stocks used in this study

| **Referred to as** | **Detailed genotype** |
|---|---|
| + | *$w^{1118}$* |
| *nanos-Gal4* | *Courtesy of Dr. Ralph Neumüller* |
| *MTD-Gal4* | *P{otu-GAL4::VP16.R}1, w*; P{GAL4-nos.NGT}40; P{GAL4::VP16-nos.UTR}CG6325MVD1* |
| *UAS-bam* | *w; CyO; p[UAS-bam:GFP, w+]54fa* |
| *GFP shRNA (I)* | *$y^1$ $sc^*$ $v^1$; P{VALIUM22-EGFP.shRNA.1}attP2* |
| *GFP shRNA (II)* | *$y^1$ $sc^*$ $v^1$; P{VALIUM20-EGFP.shRNA.4}attP2* |
| *GFP shRNA (III)* | *$y^1$ $sc^*$ $v^1$; P{VALIUM20-EGFP.shRNA.4}attP40/CyO* |
| *HexA shRNA* | *$y^1$ $sc^*$ $v^1$; P{TRiP.GL00023}attP2* |
| *Pgi shRNA* | *$y^1$ $v^1$; P{TRiP.HMC03362}attP40* |
| *Pfk shRNA* | *$y^1$ $sc^*$ $v^1$; P{TRiP.GL00298}attP2* |
| *Pyk shRNA* | *$y^1$ $sc^*$ $v^1$; P{TRiP.GL00099}attP2* |
| *Zw shRNA* | *$y^1$ $v^1$; P{TRiP.HMC03068}attP2* |
| *Pgd shRNA* | *$y^1$ $sc^*$ $v^1$; P{TRiP.HMC05959}attP40/CyO* |
| *bam shRNA* | *$y^1$ $v^1$; P{TRiP.HMS00029}attP2* |
| *$Pgd^{n39}$ $Zw^{lo2a}$* | *$Pgd^{n39}$ $pn^1$ $Zw^{lo2a}$* |
| *hs-bam;; $Fit^{81}$* | *P{hs-bam.O}18d, $w^{1118}$;; $fit^{81}$* |

**Table S2** – Composition of the diets used in this study.

The detailed composition of the stock solutions, product supplier references and methods to prepare the food can be found in [dx.doi.org/10.17504/protocols.io.heub3ew](dx.doi.org/10.17504/protocols.io.heub3ew).

| **Referred to as** | **Holidic medium (HUNTaa)** (a) | **Holidic medium (HUNTaa) - sucrose** (a) | **Holidic medium (FlyAA)** (b) | **Holidic medium (FlyAA) - sucrose** (b) | **Holidic medium (FlyAA) - amino acids** (b) |
|---|---|---|---|---|---|
| Essential amino acids | 60,51 ml | 60,51 ml | 60,51 ml (c) | 60,51 ml (c) | 0 ml (c) |
| L-isoleucine | 1,82 g | 1,82 g | 1,12 g | 1,12 g | 0 g |
| L-leucine | 1,21 g | 1,21 g | 2,03 g | 2,03 g | 0 g |
| Non-essential amino acids | 60,51 ml | 60,51 ml | 60,51 ml (c) | 60,51 ml (c) | 0 ml (c) |
| L-glutamate | 15,13 ml | 15,13 ml | 15,19 ml | 15,19 ml | 0 ml |
| L-tyrosine | 0,42 g | 0,42 g | 0,93 g | 0,93 g | 0 g |
| L-cysteine (HCl) | n/a | n/a | 6,83 ml (d) | 6,83 ml (d) | 0 ml |
| Cholesterol | 15 ml | 15 ml | 15 ml | 15 ml | 15 ml |
| $CaCl^2$ | 1 ml | 1 ml | 1 ml | 1 ml | 1 ml |
| $MgSO^4$ | 1 ml | 1 ml | 1 ml | 1 ml | 1 ml |
| $CuSO^4$ | 1 ml | 1 ml | 1 ml | 1 ml | 1 ml |
| $FeSO^4$ | 1 ml | 1 ml | 1 ml | 1 ml | 1 ml |
| $MnCl^2$ | 1 ml | 1 ml | 1 ml | 1 ml | 1 ml |
| $ZnSO^4$ | 1 ml | 1 ml | 1 ml | 1 ml | 1 ml |
| Nucleic acids & Lipids | 8 ml | 8 ml | 8 ml | 8 ml | 8 ml |
| Vitamins | 14 ml | 14 ml | 21 ml | 21 ml | 21 ml |
| Folic acid | 1 ml | 1 ml | 1 ml | 1 ml | 1 ml |
| Acetic acid buffer | 100 ml | 100 ml | 100 ml | 100 ml | 100 ml |
| Sucrose | 17,12 g | 0 g | 17,12 g | 0 g | 17,12 g |
| Agar | 20 g | 20 g | 20 g | 20 g | 20 g |
| Propionic acid | 0 ml | 0 ml | 0,6 ml | 0,6 ml | 0,6 ml |
| Nipagin | 0 ml | 0 ml | 1,5 ml | 1,5 ml | 1,5 ml |
| Milli Q $H_2O$ | Adjust to 1 L | Adjust to 1 L | Adjust to 1 L | Adjust to 1 L | Adjust to 1 L |

(a) Used in experiments shown in Figures 4A, S1D, and S2A
(b) Used in all experiments, except the ones in Figures 4A, S1D, and S2A
(c) The amino acid stock solutions have a different proportion in the HM with improved composition
(d) L-cysteine HCl was added in a separate solution only in the HM with improved amino acid composition.

| **Referred to as** | | holidic medium (HUNTaa) (b) | holidic medium (improved AA composition) (a) |
|---|---|---|---|
| Essential amino acid stock solution (1L) | L-arginine | 8 g | 26,95 g |
| | L-histidine | 10 g | 10,8 g |
| | L-lysine (HCl) | 19 g | 22,5 g |
| | L-methionine | 8 g | 9,95 g |
| | L-phenylalanine | 13 g | 16,65 g |
| | L-threonine | 20 g | 18,3 g |
| | L-tryptophan | 5 g | 5,3 g |
| | L-valine | 28 g | 19,85 g |
| Non-essential amino acid stock solution (1L) | L-alanine | 35 g | 18,2 g |
| | L-asparagine | 17 g | 17 g |
| | L-aspartic acid | 17 g | 19,35 g |
| | L-cysteine (HCl) | 0,5 g | n/a |
| | L-glutamine | 25 g | 18,55 g |
| | Glycine | 32 g | 12,7 g |
| | L-proline | 15 g | 16,15 g |
| | L-serine | 19 g | 22,8 g |
| L-glutamate stock solution (1L) | L-glutamate | 100 g | 100g |
| L-cysteine (HCl) stock solution (1L) | L-cysteine (HCl) | n/a | 50 g |

**Table S3 –** Primers for synthesis of the probes used for in *situ hybridization* and for qPCR amplification

| **Referred to as** | **Primer sequence** |
|---|---|
| *HexA* SP6 F | 5’ gactacATTTAGGTGACACTATAGAACTTCTAACGGACGAACAG 3’ |
| *HexA* T3 R | 5’ gcaacgAATTAACCCTCACTAAAGGGGCATTGGTCAGACCGAGCT 3' |
| *Pgd* SP6 F | 5’ gactacATTTAGGTGACACTATAGAAGGCAACTCGGAGTATCAGGA 3' |
| *Pgd* T3 R | 5’ gcaacgAATTAACCCTCACTAAAGGGCGTAGAAGCTTAGGGCGGTA 3' |
| *Fit* F | 5’ TTGGTGCAGGCCAGGAATAT 3' |
| *Fit* R | 5’ CACAGGGCCAGTTGACAGAGT 3' |
| *Actin42A* F | 5' CAGGCGGTGCTTTCTCTCTA 3' |
| *Actin42A* R | 5' AGCTGTAACCGCGCTCAGTA 3' |
| *RpL32* F | 5' GCCCAAGATCGTGAAGAAGC 3' |
| *RpL32* R | 5' GCACTCTGTTGTCGATACCCTTG 3' |

**Table S4** – Chemical composition of buffers used in *in situ* hybridization and carbohydrate measurement protocols

| Referred to as | Chemical composition |
|---|---|
| *2x Carbonate buffer* | 120 mM $Na_2CO_3$<br>80 mM $NaHCO_3$<br>pH to 10.2 |
| *Hybridization buffer* | 50% formamide<br>5x Saline sodium citrate<br>100 µg/mL Heparin<br>0.1% Tween 20<br>100 µg/mL sonicated and boiled salmon sperm DNA |
| *Pre-hybridization buffer* | 50% formamide<br>4x Saline sodium citrate<br>0.1% Tween20 |
| *Alkaline phosphatase (AP) buffer* | 100 mM NaCl<br>50 mM MgCl<br>100 mM Tris pH 9.5<br>0,1% Tween 20 |
| *Trehalase buffer (TB)* | 5 mM Tris pH 6.6<br>137 mM NaCl<br>2.7 mM KCl |
